# Inositol Trisphosphate Kinase and Diphosphoinositol Pentakisphosphate Kinase Enzymes Constitute the Inositol Pyrophosphate Synthesis Pathway in Plants

**DOI:** 10.1101/724914

**Authors:** Olusegun Adepoju, Sarah P. Williams, Branch Craige, Caitlin A. Cridland, Amanda K. Sharpe, Anne M. Brown, Eric Land, Imara Y. Perera, Didier Mena, Pablo Sobrado, Glenda E. Gillaspy

**Author notes:** corresponding author address: 111 Engel Hall, Department of Biochemistry, Virginia Tech, Blacksburg, VA 24061.

## Abstract

Inositol pyrophosphates (PP-InsPs) are an emerging class of “high-energy” intracellular signaling molecules containing one or two diphosphate groups attached to an inositol ring, with suggested roles in bioenergetic homeostasis and inorganic phosphate (P*i*) sensing. Information regarding the biosynthesis of these unique class of signaling molecules in plants is scarce, however the enzymes responsible for their biosynthesis in other eukaryotes have been well described. Here we report the characterization of the two Arabidopsis VIP kinase domains, a newly discovered activity of the Arabidopsis ITPK1 and ITPK2 enzymes, and the subcellular localization of the enzymes involved in the synthesis of InsP_6_ and PP-InsPs. Our data indicate that AtVIP1-KD and AtVIP2-KD act primarily as 1PP-specific Diphosphoinositol Pentakisphosphate Kinases (PPIP5) Kinases. The AtITPK enzymes, in contrast, can function as InsP_6_ kinases, and thus are the missing enzyme in the plant PP-InsP synthesis pathway. Together, these enzyme classes can function in plants to produce PP-InsPs, which have been implicated in signal transduction and P*i* sensing pathways. We measured a higher InsP_7_ level (increased InsP_7_/InsP_8_ ratio) in *vip1/vip2* double loss-of-function mutants, and an accumulation of InsP_8_ (decreased InsP_7_/InsP_8_ ratio) in the 35S:*VIP2* overexpression line relative to wild-type plants. We also report that enzymes involved in the synthesis of InsPs and PP-InsPs accumulate within the nucleus and cytoplasm of plant cells. Our work defines a molecular basis for understanding how plants synthesize PP-InsPs which is crucial for determining the roles of these signaling molecules in processes such as P*i* sensing.

**SIGNIFICANCE STATEMENT:** Inositol pyrophosphate signaling molecules are of agronomic importance as they can control complex responses to the limited nutrient, phosphate. This work fills in the missing steps in the inositol pyrophosphate synthesis pathway and points to a role for these molecules in the plant cell nucleus. This is an important advance that can help us design future strategies to increase phosphate efficiency in plants.

## INTRODUCTION

Inositol phosphates (InsPs) are part of a chemical signaling language used by most eukaryotic organisms, including plants (Gillaspy, 2011). The InsP synthesis pathway makes use of a collection of inositol phosphate kinases that function to phosphorylate specific substrates (Shears and Wang, 2019), resulting in the production of InsP signaling molecules that convey signaling information within the cell (Irvine and Schell, 2001, Hatch and York, 2010, Shears *et al*., 2012). In plants, InsPs have been studied in connection with several different developmental processes (Gillaspy, 2011), auxin and jasmonic acid pathways via binding to receptors (Tan *et al*., 2007, Sheard *et al*., 2010, Laha *et al*., 2015), and more recently in inorganic phosphate (P*i*) sensing (Kuo *et al*., 2014, Wild *et al*., 2016, Kuo *et al*., 2018).

A unique subclass of InsP signaling molecules containing one or two diphosphate groups are gaining significant attention as important players in signal transduction and regulation of cellular metabolism (Shears, 2007, Shears, 2015, Williams *et al*., 2015). These molecules are collectively named Inositol Pyrophosphates (PP-InsPs), and are considered “high energy” signaling molecules due to the considerable amount of free energy released upon hydrolysis of their pyrophosphate groups (Stephens *et al*., 1993). PP-InsPs are derived from phosphorylation of inositol hexakisphosphate (InsP_6_), one of the most abundant InsPs in the plant cell (see Figure 1) (Raboy, 2003), and a major phosphate sink in plant seeds (i.e. as phytate) (Raboy, 2007). The most common PP-InsPs contain 7 or 8 phosphates and hence are referred to as InsP_7_ and InsP_8_ (Shears, 2015).

**Figure 1.**
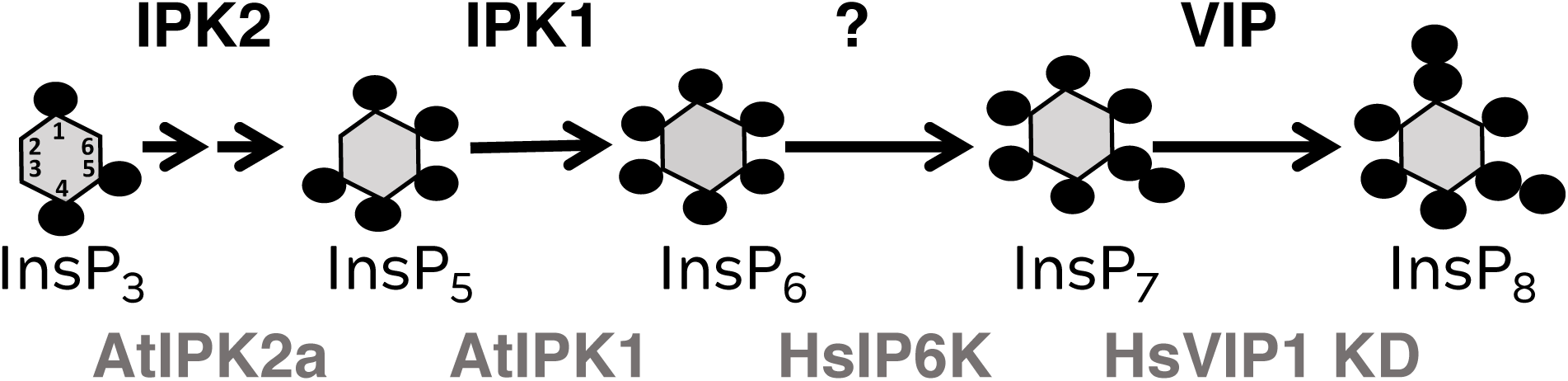
PPx-InsP Synthesis Pathway. Schematic representation of PPx-InsP synthesis pathway starting from Ins(1,4,5)P_3_. Locants are given for Ins(1,4,5)P_3_ and other molecules are similarly oriented. Enzymes (in black) on top function in the plant synthesis pathway: IPK2 = Inositol Polyphosphate Multikinase, IPK1 = Inositol Pentakisphosphate 2-Kinase, VIP = Diphosphoinositol Pentakisphosphate Kinase. The question marks denotes the lack of an identified Inositol Hexakisphosphate Kinase (IP6K) enzyme in plants. Note that the structure of InsP_7_ and InsP_8_ from plants has not yet been determined. Synthesis of substrates for this study are on the bottom in gray.

We previously showed that plants synthesize and accumulate both InsP_7_ and InsP_8_, and along with others identified the Arabidopsis *Vip/Vih* genes as capable of complementing yeast mutants defective in PP-InsP synthesis (Desai *et al*., 2014, Laha *et al*., 2015). The Arabidopsis genome contains two *Vip/Vih* genes (Desai *et al*., 2014, Laha *et al*., 2015) that encode enzymes predicted to function as Diphosphoinositol Pentakisphosphate Kinases (PPIP5Ks) (Choi *et al*., 2007). The Arabidopsis VIPs, AtVIP1 and AtVIP2, share 94% similarity at the amino acid level and 59% and 50% similarity with the human VIP1 (Fridy *et al*., 2007) and yeast VIP1 (Mulugu *et al*., 2007) respectively (Desai et al., 2014). Additionally, they share a conserved dual domain architecture housing an N-terminal ATP-grasp kinase domain and a C-terminal histidine phosphatase domain (Mulugu *et al*., 2007). VIPs phosphorylate 5PP-Ins(1,2,3,4,6)P_5_ at the 1-position, resulting in a 1-pyrophosphate group (1PP) (Lin *et al*., 2009, Wang *et al*., 2011). In other organisms, the conversion of InsP_6_ to InsP_7_ is catalyzed by InsP_6_ Kinases that form a pyrophosphate at the 5-position (5PP) (Saiardi *et al*., 1999, Saiardi *et al*., 2001, Draskovic *et al*., 2008). Since all plant genomes studied to date do not contain genes predicted to encode InsP_6_ Kinases, and the *AtVip* genes can restore mutant yeast synthesis of InsP_7_ (Desai *et al*., 2014, Laha *et al*., 2015), we previously hypothesized that the AtVIPs catalyze the two penultimate synthesis steps in the pathway, converting InsP_6_ to InsP_7_, and InsP_7_ to InsP_8_ (Desai *et al*., 2014).

Given the importance of the PP-InsPs in pathways such as P*i* sensing (Azevedo and Saiardi, 2017, Jung *et al*., 2018), we sought to characterize the biochemistry and subcellular locations of AtVIP1 and AtVIP2, and other enzymes acting in this pathway. To do this, we established a system to enzymatically produce substrates in the pathway, and used recombinant enzymes *in vitro* to show that the VIPs and another class of enzymes, the Inositol Trisphosphate Kinases (ITPKs) (Abdullah *et al*., 1992, Shears, 2009), most likely act in concert to catalyze the last two reactions in the PP-InsP synthesis pathway. InsP profiling experiments of a *vip1/vip2* double mutant and a VIP2 overexpressor (OX) line support these roles. We also investigated the subcellular distribution of many enzymes involved in the PP-InsP synthesis pathway, by using a transient expression system and confocal imaging. It has been proposed that regulation of PP-InsP synthesis may be specific for different developmental stages and/or takes place in different subcellular compartments within the cell. Our results suggest that the entire PP-InsP synthesis pathway may function in the nucleus and cytoplasm, and provides a new paradigm for understanding this critical signaling pathway.

## RESULTS

### Purification of AtVIP1-KD and AtVIP1-KD Recombinant Proteins

To determine the substrate preference of the AtVIP enzymes, we sought to express AtVIP full-length recombinant proteins in *E. coli.* As has been the case with other vertebrate VIPs, this approach was not successful in producing intact, soluble protein. We then opted to express and purify recombinant proteins corresponding to residues 1-361 for AtVIP1, and residues 1-362 for AtVIP2 as Glutathione S-Transferase (GST) fusions. These regions each contain a RimK/ATP-grasp domain, which is predicted to encode the VIP kinase domain (KD) (Fridy *et al*., 2007, Mulugu *et al*., 2007). Recombinant proteins were expressed and purified by affinity chromatography. SDS-PAGE analysis of these purified fusion proteins reveals the expected sizes of 68.6 kDa and 68.7 kDa for the AtVIP1-KD and AtVIP2-KD, respectively (Figure S1A). We also identified lower molecular weight proteins that immuno-reacted with the anti-glutathione (α-GST) antibody that we predict to be free GST or breakdown products (Figure S1B).

### Substrate Synthesis for VIP Activity Assays

To test the inositol phosphate kinase activity of recombinant AtVIP1-KD and AtVIP2-KD fusion constructs, we utilized an *in vitro* substrate synthesis system with purified enzymes and commercially available [^3^H]Ins(1,4,5)P_3_, which has been described and used by others (Fridy *et al*., 2007, Mulugu *et al*., 2007, Weaver *et al*., 2013). We expressed and purified recombinant fusion proteins with inositol phosphate kinase activities including Inositol Polyphosphate Multikinase 2 alpha (AtIPK2α) (Stevenson-Paulik *et al*., 2002), Inositol Pentakisphosphate 2-kinase (AtIPK1) (Stevenson-Paulik *et al*., 2005), the human Inositol Hexakisphosphate Kinase 1 (HsIP6K) (Saiardi *et al*., 1999), and the kinase domain of the human PPIP5K (HsVIP1-KD) (Fridy *et al*., 2007) (Figure 1). Specifically, we incubated commercial [^3^H]Ins(1,4,5)P_3_ with purified AtIPK2α, resulting in the production of [^3^H]Ins(1,3,4,5,6)P_5_ (Figure 2A). This product was incubated with AtIPK1, resulting in the formation of [^3^H]InsP_6_ (Figure 2B). For synthesis of PP-InsPs, we incubated [^3^H]InsP_6_ with HsIP6K1, to generate [^3^H]5PP-InsP_5_ (herein referred to as 5-InsP_7_) (Figure 2C). To generate [^3^H]1PP-InsP_5_ (herein referred to as 1-InsP_7_), we incubated the HsVIP1KD with [^3^H] InsP_6_. To generate [^3^H]1,5-PP_2_-InsP_4_ (InsP_8_), we used reactions containing 5-InsP_7_ with HsVIP1-KD (Figure 2D).

**Figure 2.**
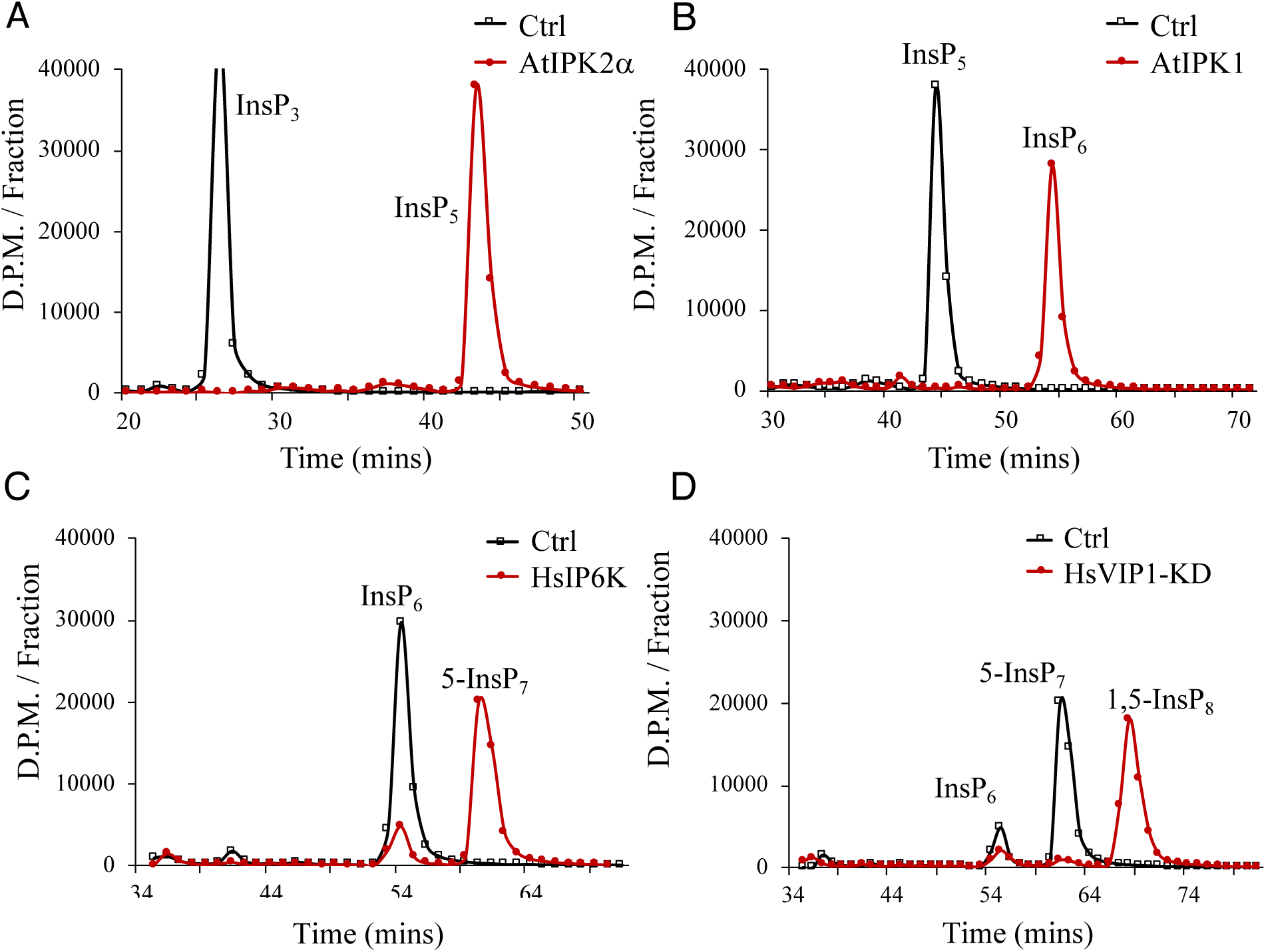
Sequential Synthesis of Substrates for Enzyme Assays. A, Incubation of 0.13 µM of commercial [^3^H]Ins(1,4,5)P_3_ with either buffer (Ctrl; open squares) or 5 µg recombinant fusion AtIPK2α (red circles) in a 100 µL reaction volume at 37°C for 60 minutes. B, Enzymatically synthesized [^3^H]Ins(1,3,4,5,6)P_5_ from panel A was incubated with either buffer (Ctrl; open squares) or 5 µg recombinant fusion AtIPK1 (red circles) for 60 minutes. C, Incubation of enzymatically synthesized [^3^H]InsP_6_ from panel B with buffer (Ctrl, open squares) or 35 μg of recombinant fusion HsIP6K1 (red circles) for 90 minutes. D, Incubation of enzymatically synthesized [^3^H]5-InsP_7_ from panel C with either buffer (Ctrl, open squares) or 7.6 μg of recombinant fusion HsVIP1-KD for 90 minutes. All reactions were terminated by heat-inactivation of enzymes at 90°C for 3 minutes followed by HPLC analysis of products. Data shown are representative of three independent syntheses.

### Arabidopsis Encoded AtVIP1-KD and AtVIP2-KD are Functional VIP/PPIP5Kinases

Given that the yeast and human VIP enzymes are thought to act primarily on 5-InsP_7_ (Choi *et al*., 2007, Fridy *et al*., 2007), we incubated both AtVIP1-KD and AtVIP2-KD with 5-InsP_7_ and found robust kinase activity (Figure 3A,B). The elution time of the product is consistent with the product being InsP_8_. We also tested whether the AtVIP-KDs could phosphorylate InsP_6_ as well as 5-InsP_7._ Under our *in vitro* assay conditions, AtVIP1-KD and AtVIP2-KD show very little kinase activity towards InsP_6_, even with an extended, 90 minute incubation (Figure 4C, 4D). We note that sometimes these extended reactions did result in 3-4% substrate conversion to InsP_7_ with both AtVIP1-KD and AtVIP2-KD.

**Figure 3.**
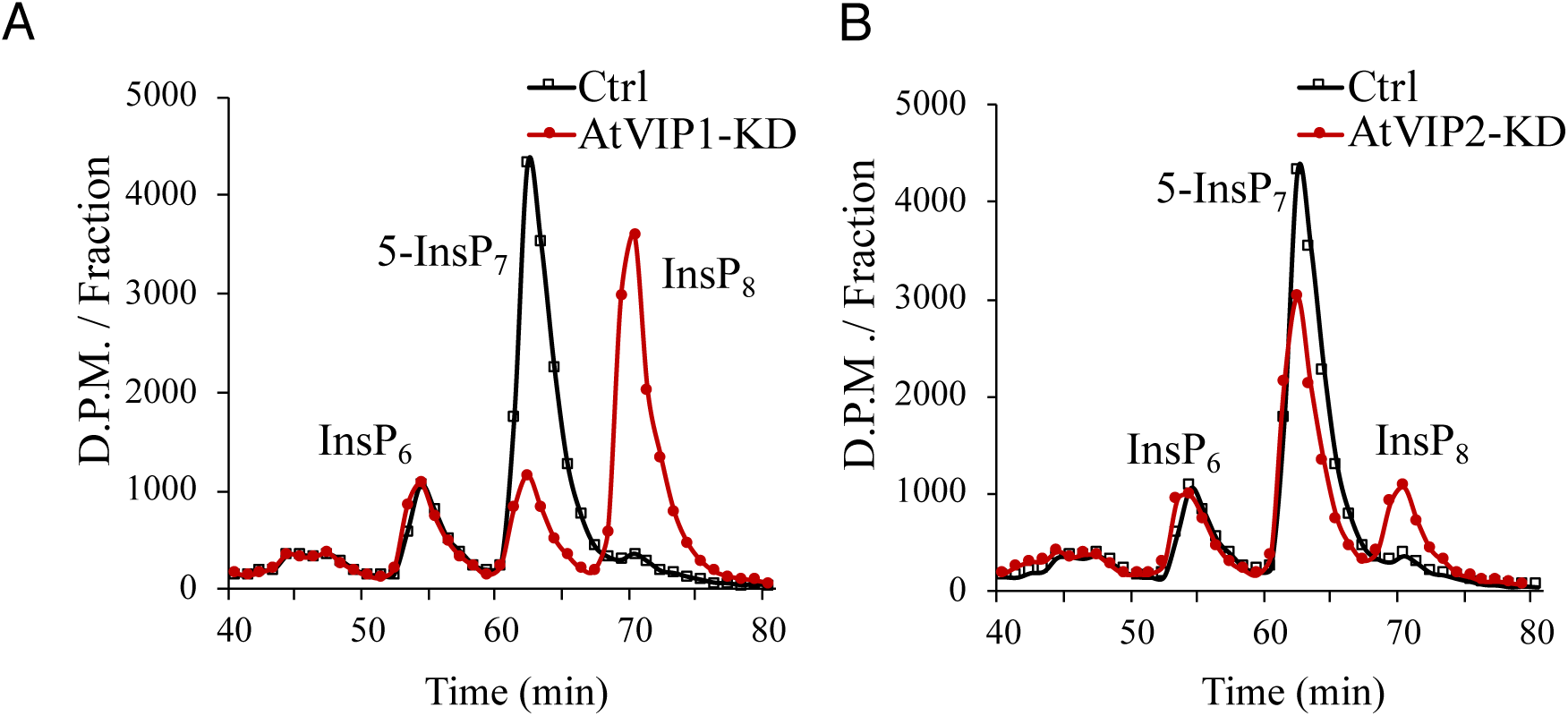
PPIP5 Kinase Activity of AtVIP1-KD and AtVIP2-KD. Enzymatically synthesized 5-InsP_7_ from 0.1 µM commercial [^3^H]InsP_6_ and recombinant HsIP6K was incubated with A, Buffer (Ctrl, open squares) or 7.6 µg of AtVIP1-KD (red circles). B, Buffer (Ctrl, open squares), or 7.6 µg of AtVIP2-KD (red circles). Reaction was incubated at 37°C for 30 minutes. All reactions were terminated with the addition of 2.5N HCl, and the reaction products were separated on an anion exchange HPLC. Data shown are representative of two independent replicates for each panel.

**Figure 4.**
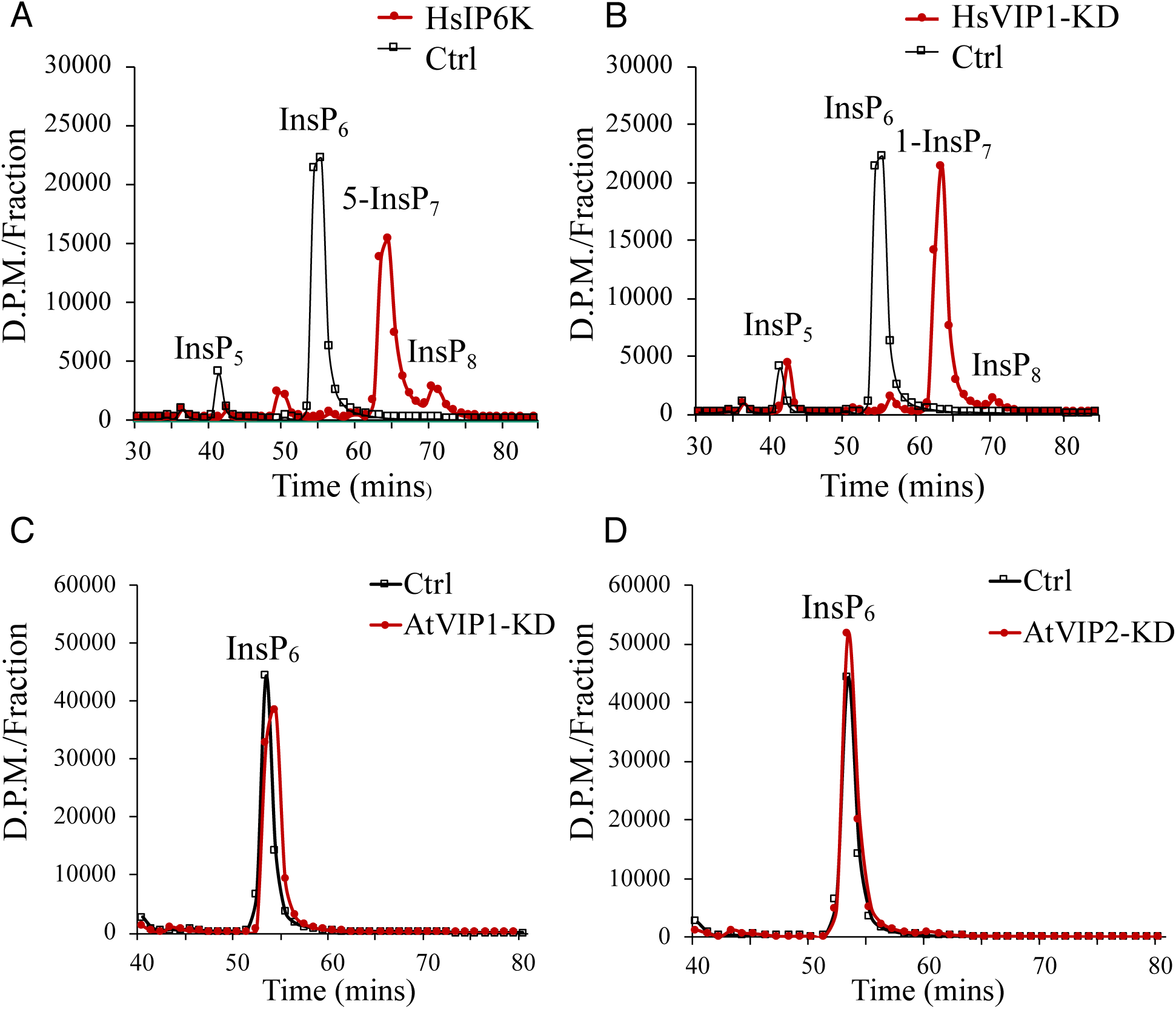
IP6Kinase Activity Assays. A, [^3^H]InsP_6_ enzymatically synthesized from 26 nM commercial [^3^H]Ins(1,4,5)P_3_ using purified recombinant AtIPK2α and AtIPK1 was incubated with either buffer (Ctrl, open squares) or 7.6 µg of purified recombinant HsIP6K (red circles). B, Buffer (Ctrl, open squares) or 7.6 µg of purified recombinant HsVIP1-KD (red circles). C, Buffer (Ctrl, open squares) or 7.6 µg of purified recombinant AtVIP1-KD (red circles). D, Buffer (Ctrl, open squares) or 7.6 µg of purified recombinant AtVIP2-KD (red circles). All reactions were incubated at 37°C for 90 minutes, and reactions were terminated by heat-inactivation of enzyme at 90°C for 3 minutes followed by HPLC analysis of products. Data shown are representative of three independent replicates.

We next tested whether 1-InsP_7_ was a substrate for purified recombinant AtVIP1-KD and AtVIP2-KD. Neither enzyme effectively converted 1-InsP_7_ to a more phosphorylated form (Figure 5 B,C), suggesting that AtVIP1-KD and AtVIP2-KD are functional enzymes that phosphorylate the 1-position of 5-InsP_7_ (5PP-InsP_5_) yielding an InsP_8_ product. We also tested whether the AtVIP-KDs could phosphorylate InsP_5_ or 5PP-InsP_4_, and found neither of these molecules are phosphorylated by the AtVIPs (Supplemental Figures S2 and S3). Together our data on substrate preference of both AtVIP-KDs indicates that their activity is similar to VIP/PPIP5 Kinases characterized from other organisms in that *in vitro*, they robustly phosphorylate 5-InsP_7_.

**Figure 5.**
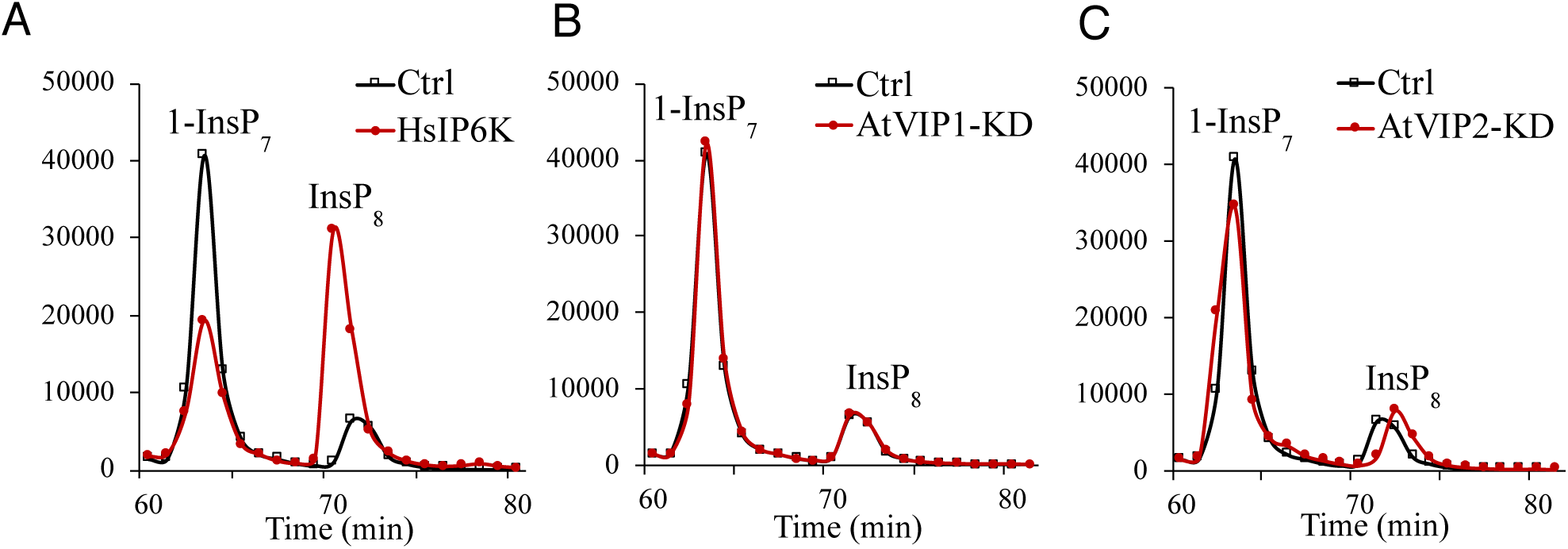
Lack of Kinase Activity of Recombinant AtVIP1-KD and AtVIP2-KD with 1-InsP_7_. Enzymatically synthesized 1-InsP_7_ from 0.1 µM of commercial [^3^H]InsP_6_ and 19 µg purified recombinant HsVIP1-KD was incubated with A, Buffer (Ctrl, open squares) or 19 µg purified recombinant HsIP6K1 (red circles). B, buffer (Ctrl, open squares) or 7.6 µg purified recombinant AtVIP1-KD (red circles). C, Buffer (Ctrl, open squares) or recombinant AtVIP2-KD (red circles). All reactions were incubated at 37°C for 90 minutes, terminated by incubation at 90°C for 3 minutes; reaction products were analyzed using an anion exchange HPLC. Data shown are representative of two independent replicates.

### Arabidopsis ITPK1 and ITPK2 have IP6K Activity

Since our characterization of the AtVIP-KDs indicates that they are not capable of generating InsP_7_, this begs the question of how InsP_7_ is synthesized in plants. Plant genomes only contain VIP homologues and are missing a bonafide InsP6K gene (Desai *et al*., 2014). Given this, we sought to identify a novel plant enzyme capable of converting InsP_6_ to InsP_7_. We reasoned that such a gene would encode a protein with known inositol phosphate 5-kinase homology. This narrowed the spectrum of our focus to the inositol polyphosphate kinase 2 enzymes which are known 6-/3-/5-kinases (Stevenson-Paulik *et al*., 2002), and the Arabidopsis inositol (1,3,4) triphosphate 5-/6-kinases (ITPKs) (Wilson and Majerus, 1997, Sweetman *et al*., 2007). There are four AtITPK genes encoded in the Arabidopsis genome (AtITPK1, AtITPK2, AtITPK3, AtITPK4) (Wilson and Majerus, 1997, Sweetman *et al*., 2007), and four characterized genes from soybean as well (Stiles *et al*., 2008). These gene products, along with a potato ITPK (Caddick *et al*., 2008), encode ATP-grasp domain containing proteins that are very structurally similar to the kinase domain of the human PPIP5K, even though they have limited sequence identity (Wang *et al*., 2011).

To test whether the AtITPKs can phosphorylate InsP_6_, we cloned and expressed AtITPK1 and AtITPK2 as recombinant 6X-histidine fusion proteins in *E. coli* (Figure S1A). 6X-His-tagged versions of AtITPK1 and AtITPK2 were incubated with [^3^H] InsP_6_, resulting in near complete conversion of substrate to a product more phosphorylated than InsP_6_ (Figure 6A, B). We next sought to test if the AtITPK1-generated InsP_7_ product could be a substrate for our purified recombinant AtVIP-KDs. We incubated the AtITPK1-generated InsP_7_ product with AtVIP1-KD, and found that the InsP_7_ could be phosphorylated to InsP_8_ (Figure 6C). Our results support a role for AtITPK1 and AtITPK2 in phosphorylating InsP_6_, and suggest that the AtITPKs function in place of the missing IP6K in plants.

**Figure 6.**
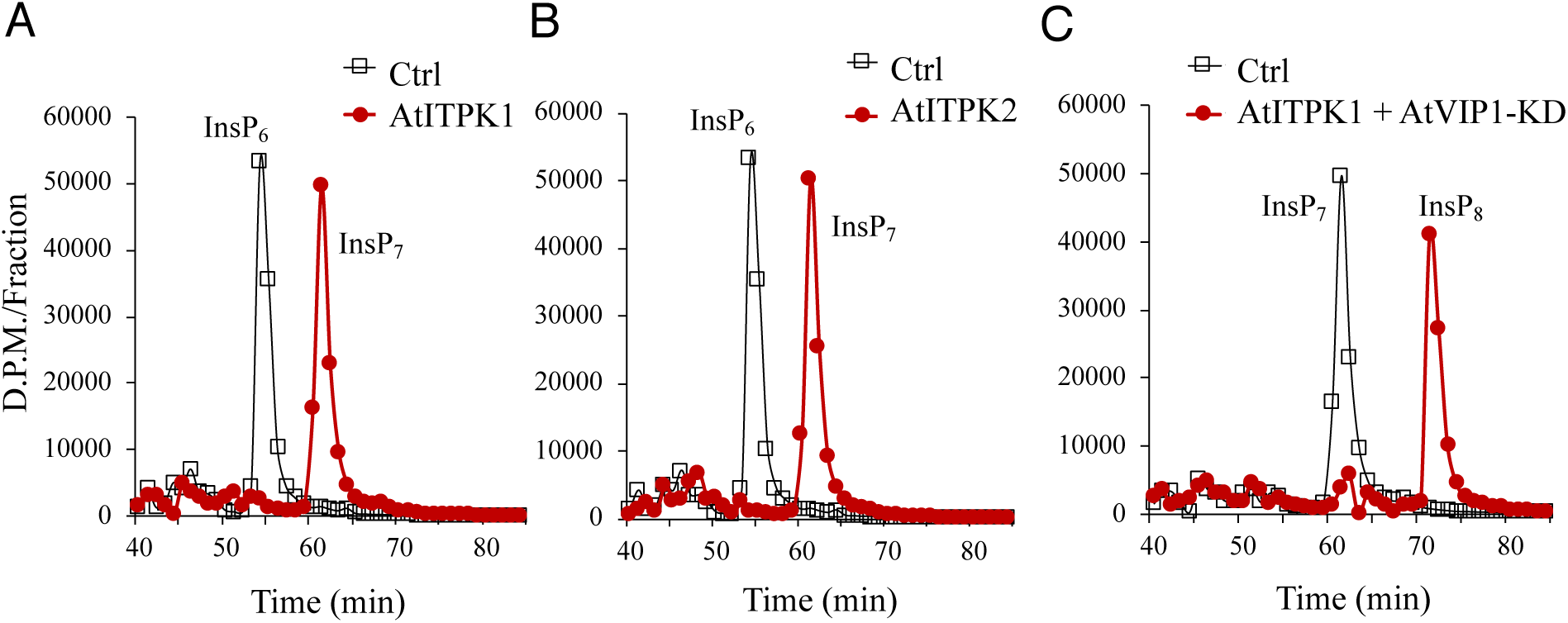
IP6Kinase Activity of AtITPK1 and AtITPK2. A, 0.1 µM commercial [^3^H]InsP_6_ was incubated with buffer (Ctrl, open squares) or 7.6 µg recombinant AtITPK1 (red circles). B, Incubation with buffer (Ctrl, open squares) or 7.6 µg recombinant AtITPK2. C, Reaction product of commercial [^3^H]InsP_6_ and AtITPK1 was incubated with buffer (Ctrl, open squares) or 7.6 µg recombinant AtVIP1-KD (closed circles) All reactions were incubated at 37°C for 90 minutes followed by termination of reaction by heat inactivation of the enzyme at 90°C for 3 minutes. Reaction products were separated by anion exchange HPLC. Data shown are representative of two independent replicates.

### *VIP* Genes Impact Inositol Pyrophosphate Accumulation

To determine if the plant *VIP* genes impact the levels of inositol pyrophosphates in plants, we isolated multiple genetic mutants in both *vip1* (*vip1-1* GK_204E06; *vip1-2* GK_008H11) and *vip2* (*vip2-1* SALK_094780*; vip2-2* SAIL_175_H09) and crossed parental lines, resulting in three unique *vip* double mutants: *vip1-1/vip2-1*, *vip1-1/vip2-2*, and *vip1-2/vip2-2*. We next performed quantitative Real-Time PCR and determined that *vip1-2/vip2-2* double mutants showed the most significant reduction in expression of *VIP1* and *VIP2* of all our double mutant lines (Figure S4). We also generated a VIP2-Green Fluorescent Protein (GFP) fusion construct driven by a 35S CaMV constitutive promoter (*VIP2 OX*), and showed that *VIP2 OX* lines accumulate an app. 146 kDa protein (Figure S4). We used [^3^H]-*myo*-inositol labelling of *vip1*-*2/vip2-2 and VIP2 OX* seedlings to quantify higher InsPs and PP-InsPs in these plants. HPLC analyses of extracts from WT, *vip1-2/vip2-2* and *VIP2 OX* plants revealed different enantiomers of InsP_4_ and InsP_5_, and a single peak corresponding to InsP_6_, InsP_7_ and InsP_8_. We found that *vip1-2/vip2-2* mutants have an elevation in InsP_7_ as compared to WT seedlings, such that the InsP_7_/InsP_8_ ratio is 140% of the WT value (Figure 7). We also determined that *VIP2 OX* plants contain an increase in InsP_8_ levels relative to *WT,* such that the InsP_7_/InsP_8_ ratio is 69% of the *WT* value (Figure 6A, B). We conclude that knock down of expression of both *VIP1* and *VIP2* genes in *vip1-2/vip2-2* mutants reduces VIP-catalyzed conversion of InsP_7_ to InsP_8_ *in vivo*. Conversely, overexpression of *AtVIP2* elevates InsP_8_ levels.

**Figure 7.**
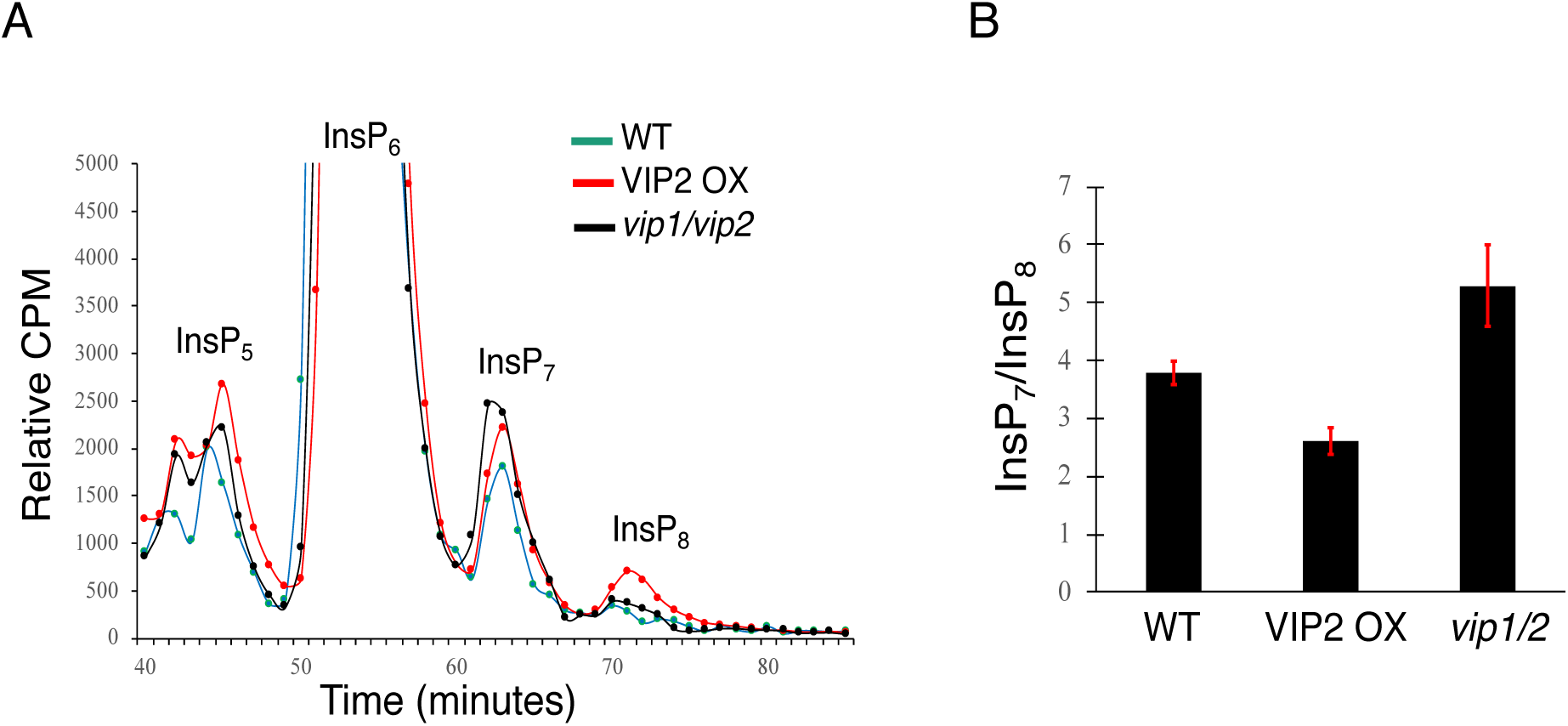
Inositol Phosphate Profiling of WT, VIP2 OX and *vip1-2/2-2* mutants. A, WT *Arabidopsis thaliana*, *AtVIP2-OE* and *vip1/vip2 (X3)* double knock-out mutant seeds were grown on semi-solid 0.5X-MS media containing 0.2% agar for 14 days and 100 μCi [^3^H]-*myo*-inositol was added for 4 days. Inositol phosphates were extracted, and analyzed on an anion exchange HPLC. B, Graph showing ratio of InsP_7_ to InsP_8_ in the different genetic constructs. The experiment was performed two times; standard error is shown.

### Localization of PP-InsP Pathway Enzymes

To determine which cellular compartments accumulate the enzymes required for PP-InsP synthesis, we constructed 35S CaMV promoter-GFP fusion proteins for each enzyme in the pathway shown in Figure 1. We transiently expressed these constructs in *N. benthamiana* leaf epidermis and assessed fusion protein integrity (Supplemental Figure S5), followed by confocal imaging to localize the fusion proteins (Figures 8-11). A time course of IPK2α-GFP, IPK2β-GFP, and IPK1-GFP expression revealed these fusion proteins accumulate in the nucleus and cytoplasm (Figure 8). All three of these fusion proteins were excluded from the nucleolus (Figure 8). The pattern of nuclear localization is consistent over a time course of 24-72 hours post infiltration, suggesting that sequential phosphorylation of Ins(1,4,5)P_3_ resulting in newly synthesized InsP_6_ could occur in the nucleus.

**Figure 8.**
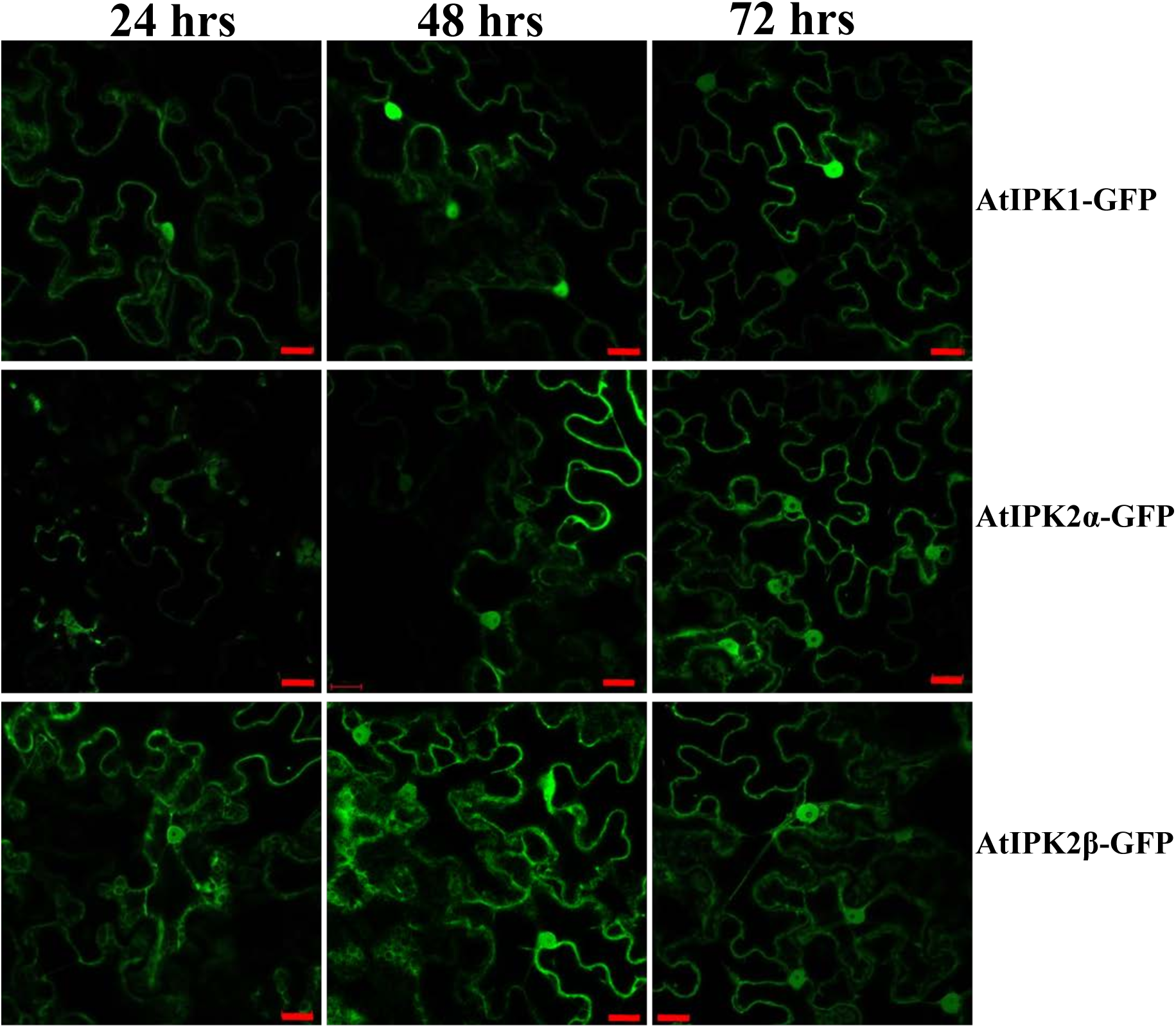
Time Course of IPK1, IPK2α, IPK2β-GFP Expression. IPK1-GFP, IPK2α-GFP, and IPK2β-GFP fusion proteins were transiently expressed in *N. benthamiana*. Leaves were imaged at 24, 48, and 72 hours post infiltration using confocal microscopy. No signal was detected at 12 hours post infiltration. Scale bar = 20 µm.

The subcellular localization of AtITPK1, AtVIP1, and AtVIP2 was assessed by transiently expressing C-terminal GFP-tag fusion proteins in tobacco leaves and confocal microscopy. Given the bipartite nature of catalytic domains in the AtVIPs, we also expressed the kinase domain (KD) with a C-terminal GFP tag, and the phosphatase domain (PD) with an N-terminal GFP tag. Cells were imaged at 12, 36, 24, 48 and 72 hours post infiltration to ensure observed patterns were not a consequence of overexpression or off-target accumulation (example time courses are shown in Supplemental Figures S6 and S7). GFP fluorescence is not detectable at 12 hours post infiltration under our conditions, and at the 72-hour time point the signal from many of the GFP constructs is beginning to decrease (Figure S7). To visualize the cytoplasm or the endoplasmic reticulum (ER), the GFP constructs were co-infiltrated with plasmids encoding unconjugated mCherry or ER-mCherry (PMID 17666025).

For all the GFP constructs investigated here, we observed extensive colocalization with mCherry and partial colocalization with ER-mCherry (Figures 9-11), indicating a major presence in the cytoplasm, and a smaller amount in the ER. Similarly, we observed localization in the nucleus for AtITPK1-GFP, AtVIP1-KD and AtVIP2-KD, but not for AtVIP1, AtVIP1-PD, AtVIP2, and AtVIP2-PD (Figures 9-11).

**Figure 9.**
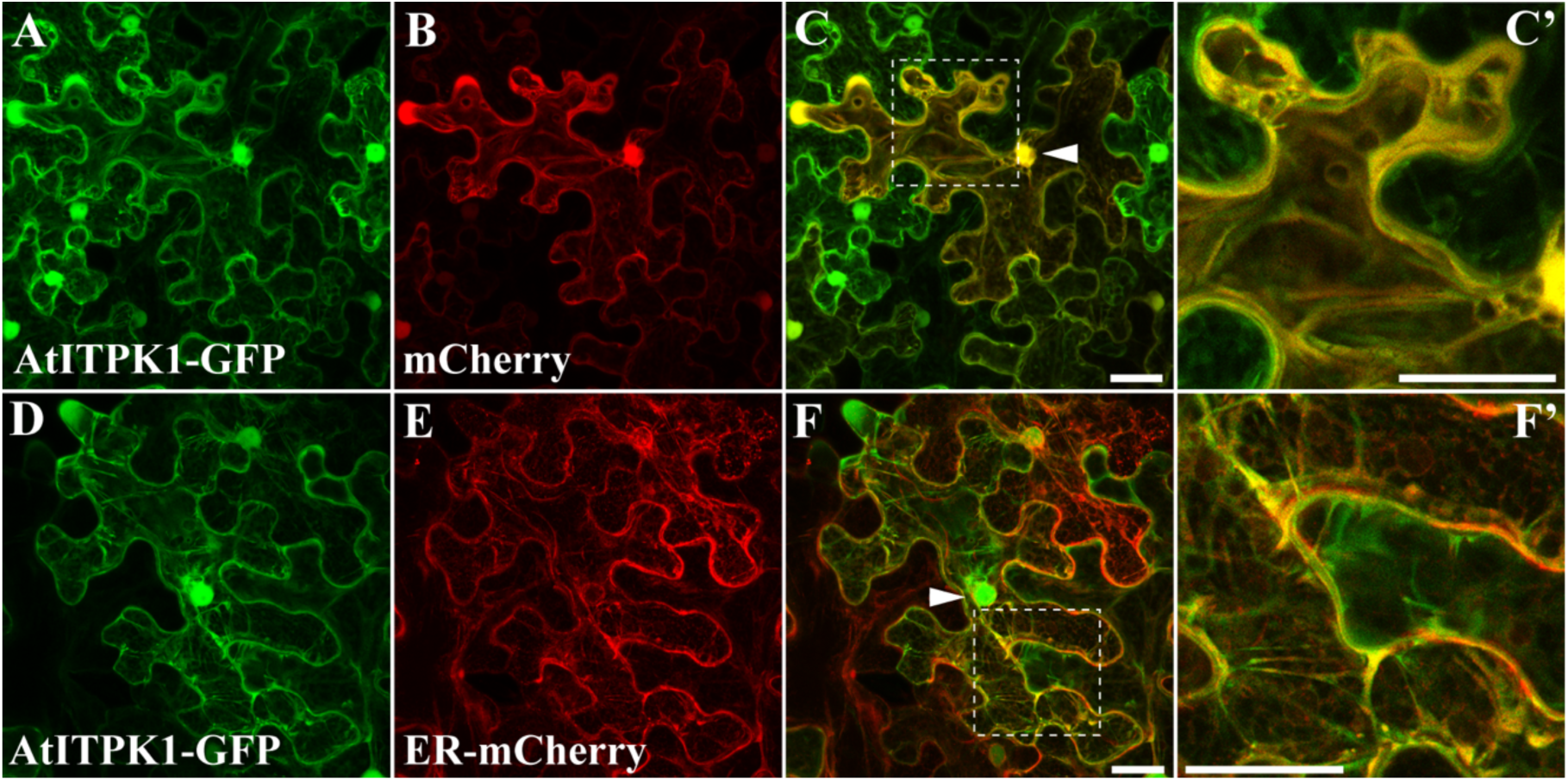
AtITPK1-GFP localization in *N. benthamiana.* *N. benthamiana* leaves were co-infiltrated with AtITPK1-GFP and mCherry (A-C’) or AtITPK1-GFP and ER-mCherry (D-F’). Z-stacks of confocal optical sections are presented as maximum intensity projections. (C’) and (F’) are 3X enlargements of the regions outlined in (C) and (F), respectively. Arrowheads mark nuclei. Scale bars = 30 µm.

**Figure 10.**
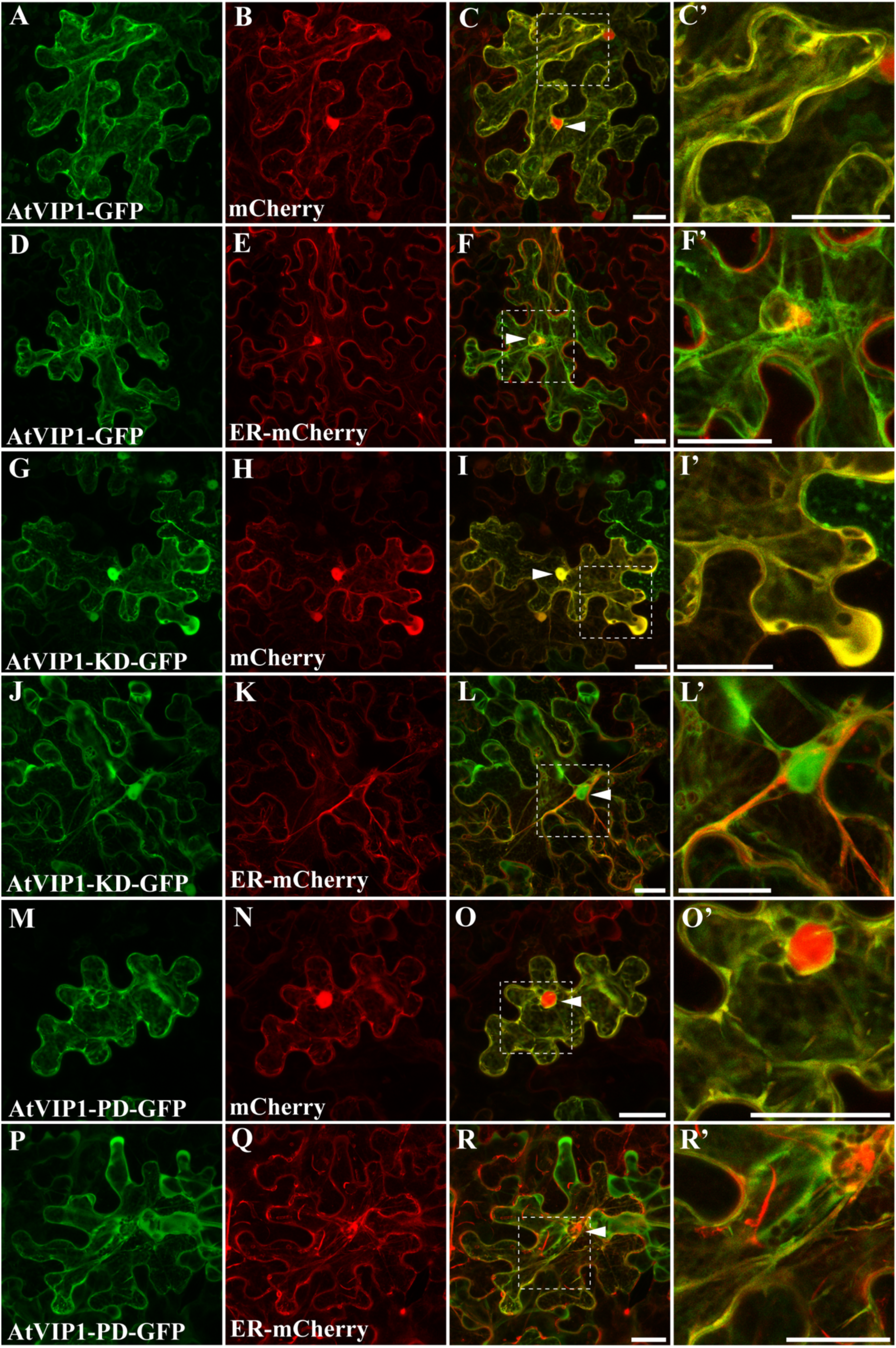
AtVIP1-GFP localization in *N. benthamiana*. *N. benthamiana* leaves were co-infiltrated with the indicated VIP1 constructs and mCherry or ER-mCherry. Z-stacks of confocal optical sections are presented as maximum intensity projections. The far-right panels are 3X enlargements of the regions outlined in the merged images. Arrowheads mark nuclei. Scale bars = 30 µm.

**Figure 11.**
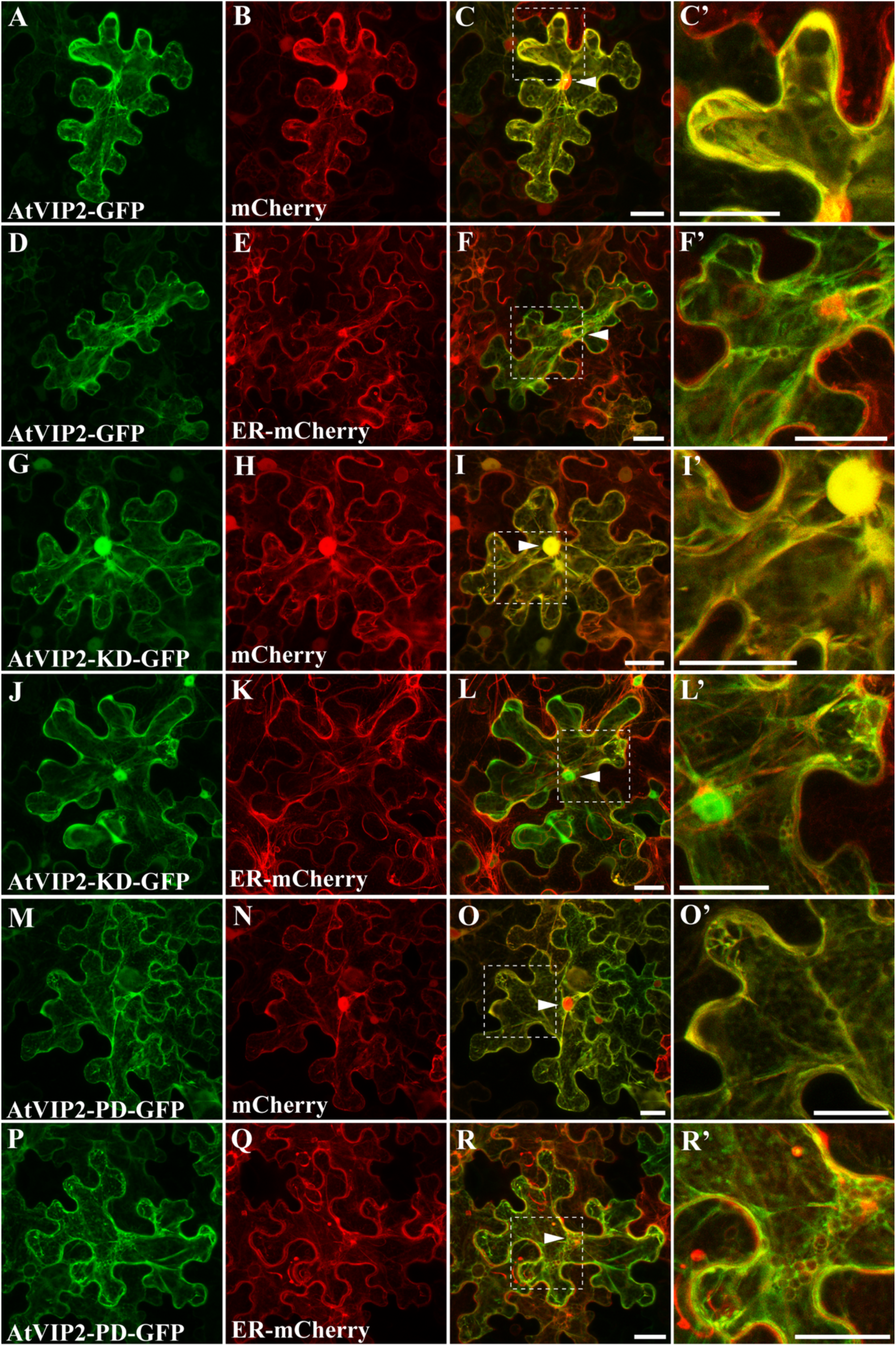
AtVIP2-GFP localization in *N. benthamiana*. *N. benthamiana* leaves were co-infiltrated with the indicated VIP2-GFP constructs and mCherry or ER-mCherry. Z-stacks of confocal optical sections are presented as maximum intensity projections. The far-right panels are 3X enlargements of the regions outlined in the merged images. Arrowheads mark nuclei. Scale bars = 30 µm.

## DISCUSSION

InsP signaling molecules have been of great interest to plant scientists since the late 1980s, starting with Ruth Satter’s work on leaflet movement and production of Ins(1,4,5)P_3_ in pulvini (Morse *et al*., 1987), to current day studies linking InsP_6_ and the PP-InsPs to P*i* sensing and homeostasis (Stevenson-Paulik *et al*., 2005, Kuo *et al*., 2014, Wild *et al*., 2016, Kuo *et al*., 2018). We have sought to fill in the missing steps in the PP-InsP synthesis pathway, and in doing this, we have defined a molecular basis for both InsP_7_ and InsP_8_ synthesis in plants (Figures 3-6). Further, we have reconstituted the synthesis pathway from Ins(1,4,5)P_3_ to InsP_8_ with plant enzymes *in vitro* (Figures 2-6), and shown that knock-down or overexpression of the AtVIPs *in vivo* results in the expected changes to InsP_7_ accumulation (Figure 7).

The ability to synthesize and separate radiolabeled substrates *in vitro* was key to work, as most of these molecules are not commercially available (Figure 2). These substrates allowed us to analyze the substrate preference of the Arabidopsis VIP kinase domains, which we show can catalyze the conversion of InsP_7_ to InsP_8_ *in vitro* (Figure 3). Our data indicates that these plant enzymes are similar in their substrate preference as compared to previously characterized yeast and mammalian enzymes, acting preferentially to phosphorylate 5-InsP_7_ substrates (Choi *et al*., 2007, Fridy *et al*., 2007, Weaver *et al*., 2013). Given that all known VIPs are 1-kinases, the most likely enantiomer of InsP_8_ in plants cells is 1,5PP_2_-InsP_4,_ which awaits further analysis by NMR for elucidation.

We coupled our *in vitro* biochemistry with an *in vivo* analysis of Arabidopsis *vip1/vip2* double mutants and AtVIP2 OX lines. While others have reported measurement of InsP_8_ in plants can be problematic (Kuo *et al*., 2018), our [^3^H]-*myo*-inositol labeling experiments allowed us to reproducibly measure accumulation of lower InsPs, along with InsP_6_, InsP_7_ and InsP_8_ in both this current study and previously (Desai *et al*., 2014). We have shown that a loss of both *VIP* genes causes a statistically significant elevation of the InsP_7_/InsP_8_ ratio, which results from elevated InsP_7_, while overexpression of *VIP2* results in a lower InsP_7_/InsP_8_ ratio (Figure 7). Together, these results support that InsP_8_ levels are maintained by the action of the *VIP* genes. Knock-out of the two human PPIP5K genes results in 2-3-fold higher 5-InsP_7_ levels in a tissue culture cell line (Gu *et al*., 2017). Thus, while our *vip1/vip2* double mutants show elevation of InsP_7_, the degree of the change is smaller and is consistent with the fact that our *vip1/vip2* double mutants are not complete nulls.

Our work has also delineated the AtITPKs as the missing enzyme in the PP-InsP pathway. Previously, two separate studies showed that the *AtVip* genes can restore InsP_7_ accumulation to yeast mutants devoid of both the IP6K and VIP enzymes (Desai *et al*., 2014, Laha *et al*., 2015). This, along with the absence of any gene encoding an IP6K homologue in plants, suggested that the AtVIP enzymes might catalyze two steps in the conversion of InsP_6_ to InsP_8_. Our biochemical analysis of the AtVIP-KDs made this seem unlikely, so we tested other Arabidopsis inositol phosphate kinases for the ability to phosphorylate InsP_6_. We determined that the Arabidopsis ITPKs can convert InsP_6_ to a more phosphorylated product (Figure 6). The ITPK family of enzymes possess an ATP-grasp fold and have been previously described as multifunctional inositol phosphate kinases capable of phosphorylating different isomers of InsP_3_ and Ins(3,4,5,6)P_4_ in plants (Wilson and Majerus, 1997, Sweetman *et al*., 2007). To our knowledge, this is the first reported *in vitro* IP6Kinase activity for any member of this family of enzymes and represents a critical piece in the puzzle of how PP-InsPs are made in plants. We suggest here that plants may have co-opted the ITPKs to perform a novel function in PP-InsP synthesis. Our observation that the reaction product of both AtITPK1 and AtITPK2 was subsequently phosphorylated by the AtVIP-KDs also suggests that the AtITPKs synthesize the 5-InsP_7_ isomer from InsP_6_, and shows that we can reconstitute the entire PP-InsP synthesis pathway *in vitro* using recombinant enzymes from plants. Given the recently suggested roles of PP-InsPs in binding to the SPX proteins that control P*i* sensing and homeostasis (Wild *et al*., 2016), as well as the finding that *itpk1* mutants are deficient in sensing P*i* (Kuo *et al*., 2018), our work suggests that lack of InsP_7_ synthesis in *itpk1* mutants may be an important factor in this mutant’s defect in P*i* sensing.

Our subcellular localization results of enzymes involved in InsP_6_, InsP_7_ and InsP_8_ synthesis suggest the presence of both nuclear and cytoplasmic locations. Specifically, we saw that the three enzymes involved in InsP_6_ synthesis have a nuclear and cytoplasmic location, as described previously by others (Xia *et al*., 2003, Kuo *et al*., 2018). Similarly, AtITPK1 and all the various AtVIP constructs displayed substantial colocalization with unconjugated mCherry, indicating localization in the cytoplasm. We also observed, to a lesser extent, partial colocalization with ER-mCherry. AtITPK1-GFP, and the kinase domains (KD) of AtVIP1 and AtVIP2, were also observed in the nucleus, but the full-length and phosphatase domains (PD) of AtVIP1 and AtVIP2 were seen to be excluded from the nucleus. This suggests that the AtVIP PD might contain a nuclear export signal (NES), since the AtVIP-KD constructs were able to access the nucleus in the absence of the PD. We note that a nuclear localization sequence (NLS) has been identified near the C-terminus of the human PPIP5K2, and mutation of this NLS increases the portion of PPIP5K2 that accumulates in the cytoplasm of mammalian cells (Yong *et al*., 2015). Similarly, the human IPK2 paralogue (IMPK) contains an NLS, along with an NES, and together these allow for nucleocytoplasmic shuttling of the enzyme in human cells (Meyer *et al*., 2012). It will be interesting to further dissect the mechanisms regulating AtVIP localization, and it is tempting to speculate that AtVIP access to the nucleus could be dynamically regulated during plant development and/or nutrient demand, resulting in a nuclear pool of InsP_8_.

This work furthers our understanding of critical biochemical and cell biological aspects of the PP-InsP synthesis pathway in plants. Given the importance of seed InsP_6_ and P*i* in agriculture, delineation of the PP-InsP synthesis pathway has important ramifications for future approaches to control P*i* sensing in plants.

## EXPERIMENTAL PROCEDURES

### Materials

[^3^H]-*myo*-Inositol (20 Ci/mmol), [^3^H]Ins(1,4,5)P_3_ (17.1 Ci/mmol) and [^3^H]InsP_6_ (19.3Ci/mmol) were purchased from American Radiolabeled Chemicals (ARC), St. Louis, MO, USA. [^3^H]Ins(1,3,4,5,6)P_5_, and all other radiolabeled PP-InsPs (5PP-InsP_4_, 1PP-InsP_5_, 5PP-InsP_5_ and 1,5(PP)_2_-InsP_4_) were synthesized enzymatically with the appropriate purified recombinant enzymes.

### Plant Materials and Growth Conditions

Seeds of T-DNA mutants of *Arabidopsis thaliana* (ecotype Col-0) were obtained from the Arabidopsis Biological Resource Center (ABRC, Columbus, OH, USA). The T-DNA lines used in this study are as follows: *vip1-1* (GK_204E06), *vip1-2* (GK_008H11), *vip2-1* (SALK_094780) and *vip2-2* (SAIL_175_H09). Homozygous lines were identified by PCR using T-DNA left and right border primers and gene-specific sense or antisense primers (Supplemental Table 1). Double loss-of function mutant lines were generated by crossing *vip1-1* with *vip2-1*, *vip1-1* crossed with *vip2-1*, and *vip1-2* was crossed with *vip2-2*. All lines analyzed in this study, including Col-0 plants, were grown in parallel under identical conditions on soil (16 h light and 8 h dark, day/night temperature 23/21°C and 120 μE light intensity), and seeds of the respective last progenies were used for all analyses described. For growth in sterile conditions, seeds were sterilized in 30% (v/v) Clorox for 5 min and washed three times in ddH_2_O. Sterilized seeds were plated onto 0.5X MS media supplemented with 0.2% agar and stratified for 3 days at 4°C, grown under conditions of 16 h light (23°C) and 8 h dark (21°C) and 120 μE light intensity.

### Synthesis of InsPs and PP-InsPs

[^3^H]Ins(1,3,4,5,6)P_5_ was prepared by incubating [^3^H]Ins(1,4,5)P_3_ with recombinant AtIPK2α at 37°C for 90 minutes. Enzymatically synthesized [^3^H]Ins(1,2,3,4,5,6)P_6_ was prepared by incubating [^3^H]Ins(1,3,4,5,6)P_5_ with recombinant AtIPK1 at 30°C for 60 minutes. [^3^H]5PP-Ins(1,2,3,4,6)P_5_ and [^3^H]1PP-Ins(2,3,4,5,6)P_5_ were generated by incubating [^3^H]Ins(1,2,3,4,5,6)P_6_ at 37°C for 60 minutes and 240 minutes with recombinant human InsP_6_ Kinase (HsIP6K) and the human VIP1 Kinase Domain (HsVIP1-KD) respectively. [^3^H]5PP_2_-Ins(1,3,4,6)P_4_ was synthesized by incubating [^3^H]Ins(1,3,4,5,6)P_5_ with recombinant HsIP6K at 37°C for 60 minutes. Reaction products were separated on an anion exchange HPLC, fractions were taken and analyzed by scintillation counting, and identity of the products was determined based on their elution profile, and elution of standards in separate runs.

### Construction of Plasmids

Expression constructs and oligonucleotide primers used are summarized in Supplemental Table 1. Additional expression constructs were provided by Ryan Irving and John York (Vanderbilt University); His-AtIPK2*α* (JYB897), GST-AtIPK1 (JYB308), GST-HsIP6K (JYB 1629) and GST-HsVIP1-KD (JYB1556). Recombinant AtVIP1-KD, AtVIP2-KD, AtITPK1 and AtITPK2 were generated as follows: Total RNA was extracted from young *Arabidopsis thaliana* seedlings using the QIAGEN RNeasy Mini Kit (QIAGEN, Hilden, Germany). The RNA was reverse transcribed using the Bio-Rad iScripts reverse transcriptase (Hercules, CA, USA) to generate cDNA and primers were designed against the coding sequence of the kinase domains of AtVIP1 (At3G01310), AtVIP2 (At5G15070), and full length AtITPK1 (At5G16760) and AtITPK2 (At4G33770) with the addition of a 5’-CACC sequence for site-specific cloning into the pENTR Gateway entry vector (Thermo Fisher Scientific, Hampton, NH, USA). The PCR amplified products were gel purified using a QIAGEN DNA gel purification kit (QIAGEN, Hilden, Germany), and the purified products were cloned into the Gateway pENTR/D Topo cloning vector (Thermo Fisher Scientific, Hampton, NH, USA). and transformed into TOP10-competent *E. coli* cells (Fisher Scientific, Hampton, NH, USA). Positive entry clones were sub-cloned into a Gateway-compatible destination vector. pDEST-17 carrying an N-terminal 6X-His purification tag, or pDEST-24 carrying a C-terminal GST purification tag (Thermo Fisher Scientific, Hampton, NH, USA).

### Expression and Purification of Recombinant GST-tagged Proteins

Recombinant protein for all GST-tagged constructs (HsVIP1-KD, HsIP6K, AtIPK1, AtVIP1-KD and AtVIP2-KD) were expressed in BL21 DE3 Turbo competent *E. coli* cells (Gelantis Biotechnology, San Diego, CA, USA) through auto-induction by culturing cells in a highly-enriched terrific broth (TB) media at 18°C for 16 hours. Cells were harvested by centrifugation at 4000 x g for 20 minutes and resuspended for 30 minutes at 4° C in lysis buffer (25 mM Tris (pH 8.0), 350 mM NaCl, 1mM DTT) supplemented with 100 µg/mL lysozyme and 1 tablet of EDTA-free complete protease inhibitor (Sigma-Aldrich, St. Louis, MO, USA). Resuspended cells were lysed by sonication, centrifuged at 17,500 RPM for 60 minutes and the clarified lysate loaded onto a 5 mL GSTrap HP (GE Healthcare, Chicago, IL, USA). The GSTrap HP column was washed with wash buffer (25 mM Tris (pH 8.0), 350 mM NaCl, 1 mM DTT, 0.1 mM reduced glutathione), and protein was eluted with elution buffer (25 mM Tris (pH 8.0), 350 mM NaCl, 1 mM DTT, 20 mM reduced glutathione) at room temperature. Three elutions were combined, and proteins were concentrated using Amicon ultracentrifugal 30-kDa cut-off filters for 30 min at 4000 X g at 4°C (Eppendorf 5810 R centrifuge). The purified recombinant protein was stored in storage buffer (20 mM HEPES (pH 7.5), 150 mM NaCl, 1 mM DTT, and 25% glycerol) at 80°C. Aliquots were examined on a 10% SDS-PAGE followed by Coomassie Blue staining or Western Blot analysis with rabbit α-GST-HRP antibody (Life Technologies, Carlsbad, CA) at a 1:50,000 dilution in Tris-buffered saline (20 mM Tris, pH 7.5, 150 mM NaCl), 2.5% non-fat milk and 0.1% Tween 20, and the proteins were compared with a Low range Prestained protein standard (Bio-Rad Laboratories, Hercules, CA).

### Expression and Purification of Recombinant 6X-His-tagged Proteins

All 6X-His-tagged plasmid constructs (AtIPK2α, AtITPK1 and AtITPK2) were expressed in BL21 DE3 Turbo competent *E. coli* cells (Gelantis Biotechnology, San Diego, CA, USA) through auto-induction by culturing cells in a highly-enriched terrific broth (TB) media at 18°C for 16 hours. Cells were harvested by centrifugation at 4000 x g for 20 minutes and resuspended for 30 minutes at 4 °C in lysis buffer (25mM HEPES (pH 7.5), 350 mM NaCl, 1 mM DTT, 1 mM PMSF) supplemented with 100 µg/mL lysozyme and 1 tablet of EDTA-free complete protease inhibitor (Sigma-Aldrich, St. Louis, MO, USA). Resuspended cells were lysed by sonication, centrifuged at 17,500 RPM for 60 minutes and the clarified lysate loaded onto a 5 mL HisTrap HP (GE Healthcare, Chicago, IL, USA). The HisTrap HP column was washed with wash buffer (25 mM HEPES (pH 7.5), 350 mM NaCl, 1 mM DTT, 20 mM imidazole), and eluted with elution buffer (25 mM HEPES (pH 7.5), 350 mM NaCl, 1 mM DTT, 250 mM imidazole) at room temperature. Three elutions were combined, and proteins were concentrated using Amicon ultracentrifugal 30-kDa cut-off filters for 30 min at 4000 x g at 4°C (Eppendorf 5810 R centrifuge). The purified recombinant protein was stored in storage buffer (20 mM HEPES pH 7.5, 150 mM NaCl, 1mM DTT, and 25% glycerol) at −80°C. Aliquots were analyzed on a 10% SDS-PAGE followed by Coomassie Blue staining or Western blot analysis with goat α-6X-His-HRP antibody (Life Technologies, Carlsbad, CA) at a 1:5,000 dilution in Tris-buffered saline (20 mM Tris, pH 7.5, 150 mM NaCl), 2.5% non-fat milk and 0.1% Tween 20, and the proteins were compared with a Low range Pre-stained protein standard (Bio-Rad Laboratories, Hercules, CA). The purified recombinant protein was stored in storage buffer (20 mM HEPES (pH 7.5), 150 mM NaCl, 1mM DTT, and 25% glycerol) at −80°C.

### Inositol Phosphate Kinase Enzyme Assays

VIP kinase activity assays were performed according to (Volgmaier *et al.,* 1996) with the following modifications: 100 μL reactions containing 20 mM HEPES pH 6.2, 50 mM NaCl, 5 mM Na_2_ATP, 1 mM DTT, 6 mM MgSO4, 6 mM phosphocreatine, 24 Units/mL phosphocreatine kinase (Calbiochem, Burlington, MA, USA) and reactions were incubated at 37°C for the times indicated. Both AtVIP1-KD and AtVIP2-KD acted on 5-InsP_7_ and had linear accumulation of InsP_8_ product over the first 10 minutes of the reaction. In some cases, incubation times of up to 90 minutes were used to assess whether molecules other than 5-InsP_7_ could be acted upon by these enzymes. ITPK assays were performed in a 100 uL reaction buffer containing 20 mM HEPES pH 6.2, 50 mM NaCl, 5 mM Na_2_ATP, 1 mM DTT, 6 mM MgSO_4_, 6 mM phosphocreatine, 24 Units/mL phosphocreatine kinase (Calbiochem, Burlington, MA, USA). Reactions were incubated at 37°C for 90 minutes unless otherwise noted, and enzyme reactions were heat-inactivated at 90°C for 3 minutes. Reaction products from enzyme assays were analyzed by HPLC using a 125 X 4.6 mm Partisphere SAX-column (Sigma-Aldrich, St. Louis, MO, USA), and a previously described elution gradient (Azevedo *et al.,* 2006).

### Cloning of GFP constructs

The coding regions of IPK1, IPK2α, IPK2β, VIP1-FL, VIP1-KD, VIP1-PD, VIP2-FL, VIP2-KD, VIP2-PD, ITPK1 and ITPK2 were amplified from plasmids, WT Arabidopsis genomic or cDNA using the primers indicated in Supplemental Table 2. The PCR product corresponding to the full length coding sequence without the stop codon was cloned into pENTR™\D-TOPO® (Invitrogen Corp., Carlsbad, CA) and sequenced before recombining either VIP2PD into pK7WGF2 (N-terminus GFP) or all others into pK7FWG2 (C-Terminus GFP) (Karimi *et al*. 2002) containing the 35S Cauliflower Mosaic virus promoter and eGFP gene using the Gateway® LR Clonase™ II kit (Invitrogen Corp., Carlsbad, CA). A single colony was used to amplify the [Gene-pK7FW2/pK7WGF2] plasmid and it was sequenced before transforming Agrobacterium (strain GV3101) for transient expression in *N. benthamiana*. Cloning and expression plasmids were amplified in *E. coli* One Shot®Top10 cells (Invitrogen Corp., Carlsbad, CA) and purified from an overnight culture using DNA Mini prep kit (Qiagen Co., Valencia, CA).

### GFP Localization and Imaging

*N. benthamiana* plants were agro-infiltrated as previously described (Kapila *et al*., 1997). Briefly, Agrobacterium cultures were grown over night in liquid media. Cells were pelleted and suspended in MMA (1x MS, 10 mM MES, 200 µM acetosyringone) to an optical density of 1.0, A600 nm. Cultures were allowed to incubate at room temperature 2-4 hours before infiltration. *N. benthamiana* plants were grown under long day conditions (16 hours light) and 150 µE light. Leaf sections were imaged 12, 24, 36, 48, or 72 hours post infiltration as indicated using a Zeiss LSM 880 (Carl Zeiss). Slides were examined with a 40x C-Apochromat water immersion lens. A set of mCherry tagged organelle markers were used for co-localization experiments (Nelson *et al*., 2003). GFP was excited using a 488-nm argon laser and its fluorescence was detected at 500- to 550-nm. mCherry was imaged using excitation with a 594-nm laser and fluorescence was detected at 600-650-nm. Chlorophyll signal was collected using a 594-nm laser and emission above 650 nm was collected.

### Protein Blot Analyses of GFP-Fusion Proteins

Conditions have been previously reported (Burnette *et al*., 2003). Briefly, tissues were ground in liquid nitrogen, homogenized, cellular debris was pelleted, and SDS-bromophenol blue loading dye added to the supernatant. The supernatant was boiled, centrifuged, and the subsequent supernatant was loaded onto a polyacrylamide gel for separation. Equal amounts of protein were loaded onto gels. SDS-PAGE was followed by western blotting with a 1:10,000 dilution of rabbit anti-GFP antibody (Invitrogen Molecular Probes, Eugene, OR). A secondary goat anti-rabbit horseradish peroxidase antibody (Bio-Rad Laboratories, Hercules, CA) was used at a 1:2,500 dilution For Supplemental Figure S4, proteins were transferred to PVDF membrane and blotted proteins were incubated in a 1:10,000 dilution of mouse monoclonal anti-GFP antibody (Living Colors JL-8, Clontech) followed by 1:5000 horseradish peroxidase conjugated sheep anti-mouse IgG antibody (Amersham, Buckinghamshire, UK). Immunoreactive bands were detected using an ECL™ Prime Western Blotting Detection Reagent (Amersham, Buckinghamshire, UK). Ponceau S staining of blots was performed to ensure that equal amounts of protein within extracts were analyzed.

#### RNA preparation and qRT-PCR

RNA was isolated from 2 week-old seedlings grown on MS medium with 1% sucrose. RNA was isolated using the Plant RNeasy kit (Qiagen) and was DNAse treated with DNA-free Turbo (Invitrogen). Reverse transcription was carried out using 2 µg of total RNA Multiscribe Transcriptase (Applied Biosystems). The equivalent of 20 ng of cDNA was used for quantitative PCR using gene specific primers for *AtVip1* and *AtVip2* and SYBRgreen as described in Desai *et al*., 2014. Reactions were carried out in triplicate with PP2A as the reference gene. Relative expression was calculated by the ΔΔCT method.

## ACCESSION NUMBERS

AtIPK2*α*: At5G07370; AtIPK2β: At5G61760; AtIPK1: At5G42810; AtITPK2: At4G33770; AtVIP1: At3G01310; AtVIP2: At5G15070.

## ACKNOWLEDGMENTS

The authors would like to acknowledge John York and Ryan Irving who graciously supplied plasmids and purified proteins, Joonho Park for help with construction of GFP plasmids, Janet Donahue for expert assistance, and members of the Gillaspy lab for critical reading and comments of this manuscript. This research was supported by a National Science Foundation grant to GG, PS and IYP.

---

**Supplemental Figure S1.**
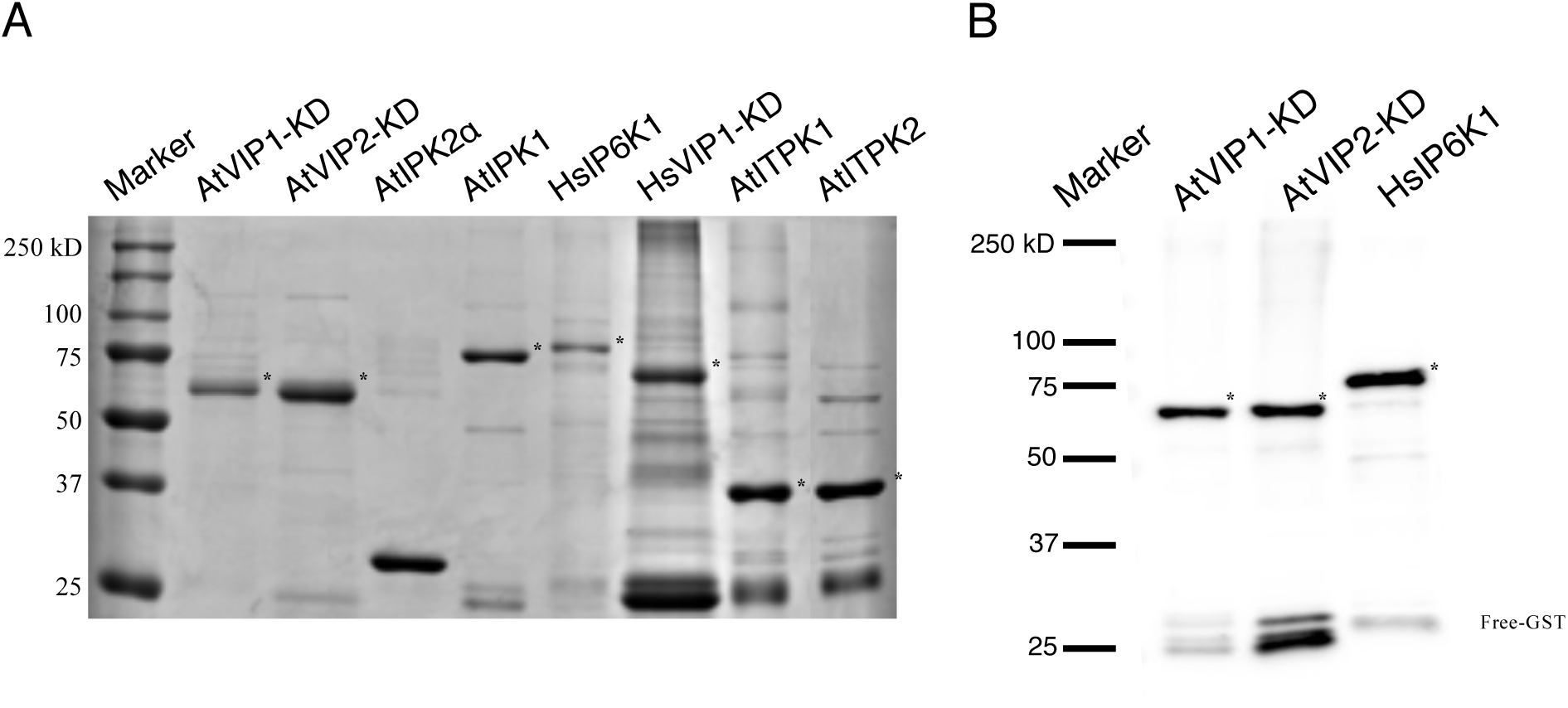
SDS-PAGE and Western Blot of Purified Recombinant Fusion Proteins Used in the Study. A, Recombinant fusion proteins were analyzed by SDS-PAGE, and stained with Coomassie blue. All target recombinant fusion proteins used in study are marked with asterisks. Recombinant GST-tagged AtVIP1-KD and AtVIP2-KD ran slightly below their expected molecular weight of 73kD, 6His-tagged AtIPK2α ran slightly below its expected molecular weight of 33kD, recombinant GST-tagged AtIPK1 was seen around 76kD, recombinant GST-tagged HsIP6K ran at the expected 75kD, recombinant GST-tagged HsVIP1-KD migrated at the expected 70kD and recombinant 6His-tagged AtITPK1 and AtITPK2 both ran slightly below their expected 36kD and 41kD respectively. B, Immunoblot analyses showed that the contaminating band around 25kD on the GST-tagged proteins is free GST as it immunoreacted with α-GST antibody.

**Supplemental Figure S2.**
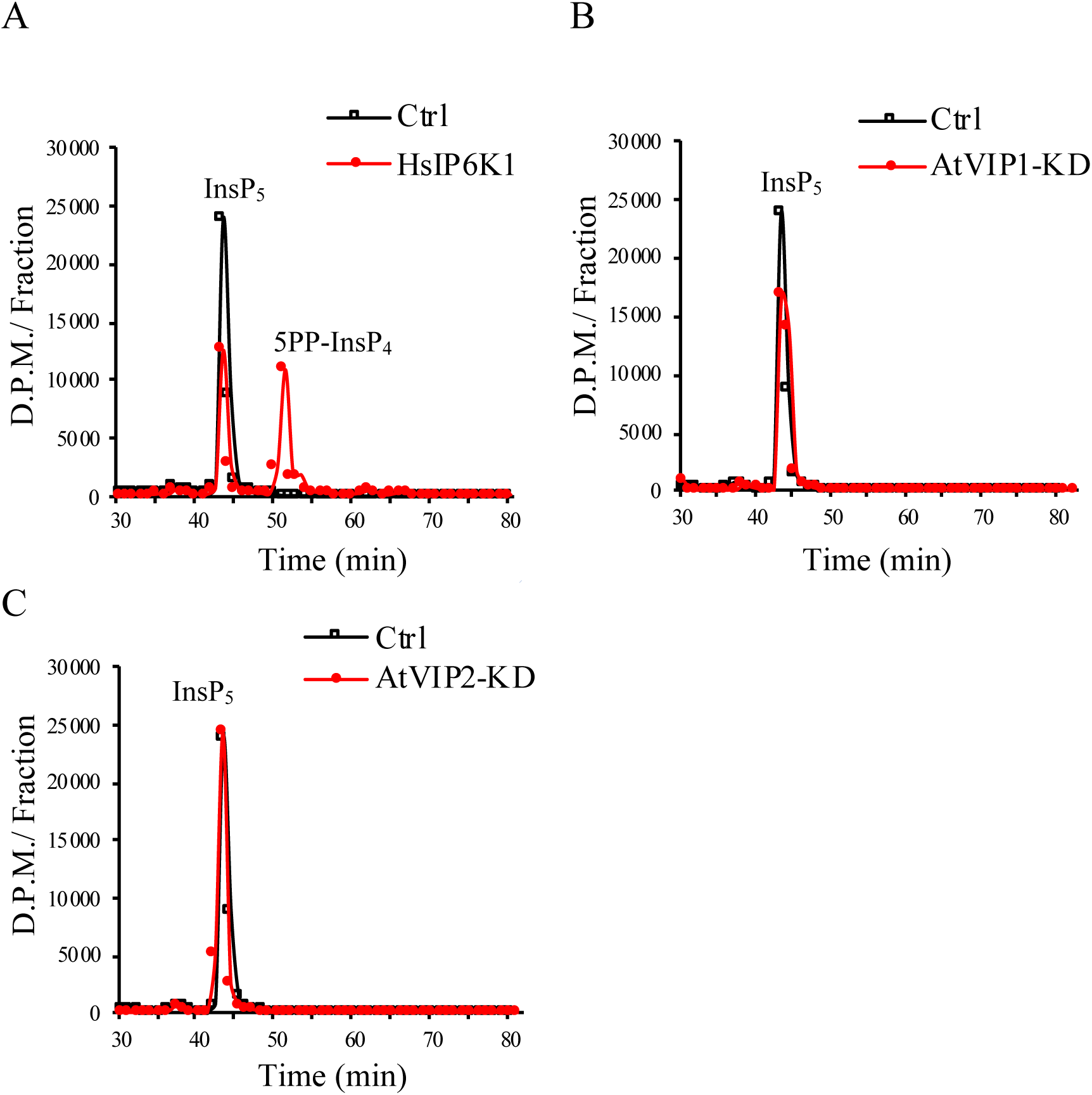
InsP5 is not a Substrate for Recombinant AtVIP1-KD or AtVIP2-KD. [^3^H]Ins(1,3,4,5,6)P_5_ enzymatically synthesized from 0.13 µM commercial [^3^H]Ins(1,4,5)P_3_ using purified recombinant AtIPK2α was incubated with A, either buffer (Ctrl, open squares) or 7.6 µg of purified recombinant HsIP6K1 (red circles). B, Buffer (Ctrl, open squares) or 7.6 µg of purified recombinant AtVIP1-KD (closed circles). C, Buffer (Ctrl, open squares) or 7.6 µg of purified recombinant AtVIP2-KD (closed circles). All reactions were incubated at 37°C for 90 minutes, and reactions were terminated by heat-inactivation of enzyme at 90°C for 3 minutes followed by HPLC analysis of products. Data shown are representative of two replicate

**Supplemental Figure S3.**
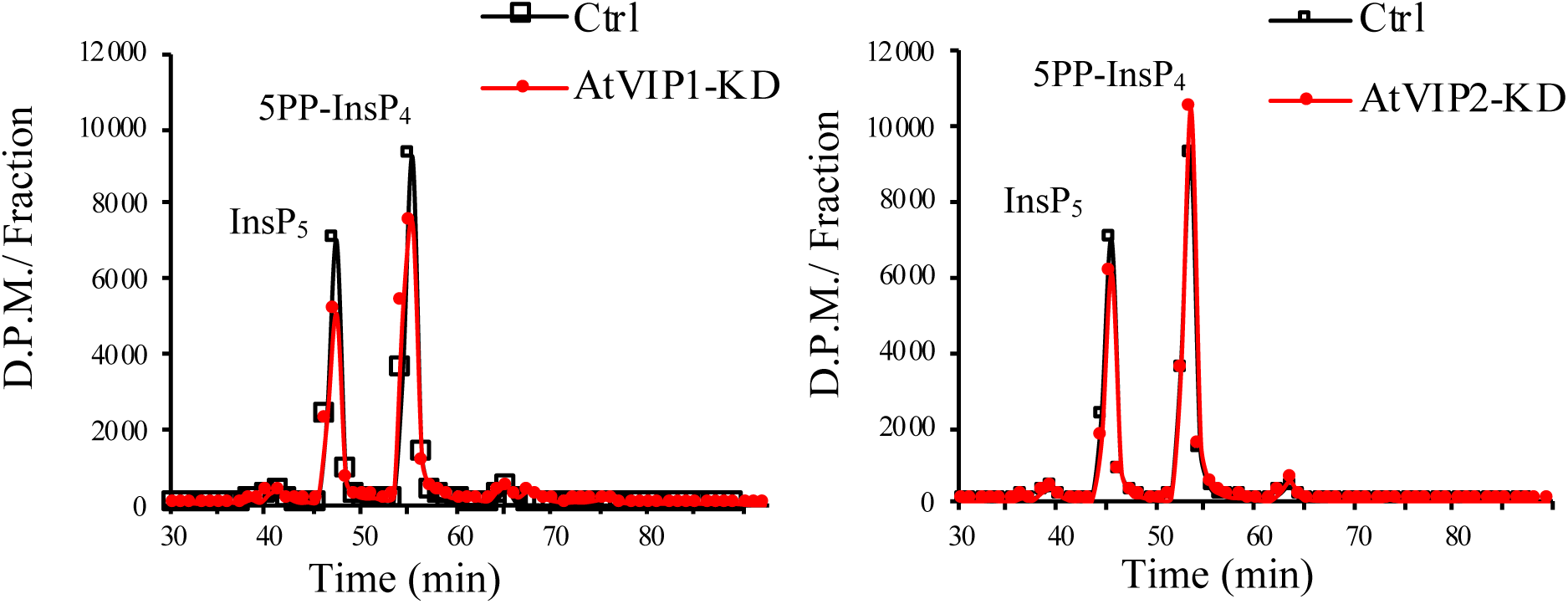
5PP-InsP_4_ is not a Substrate for Recombinant AtVIP1-KD or AtVIP2-KD. Enzymatically synthesized 5PP-InsP_4_ from [^3^H]Ins (1,3,4,5,6)P_5_ and 7.6 µg purified recombinant HsIP6K was incubated with buffer (Ctrl, open squares) or 7.6 µg purified recombinant AtVIP1-KD or AtVIP2-KD (red circles). Reactions were incubated at 37°C for 90 minutes, reactions were terminated by incubation at 90°C for 3 minutes, and reaction products were analyzed using anion exchange HPLC. Data shown are representative of two replicate experiments.

**Supplemental Figure S4.**
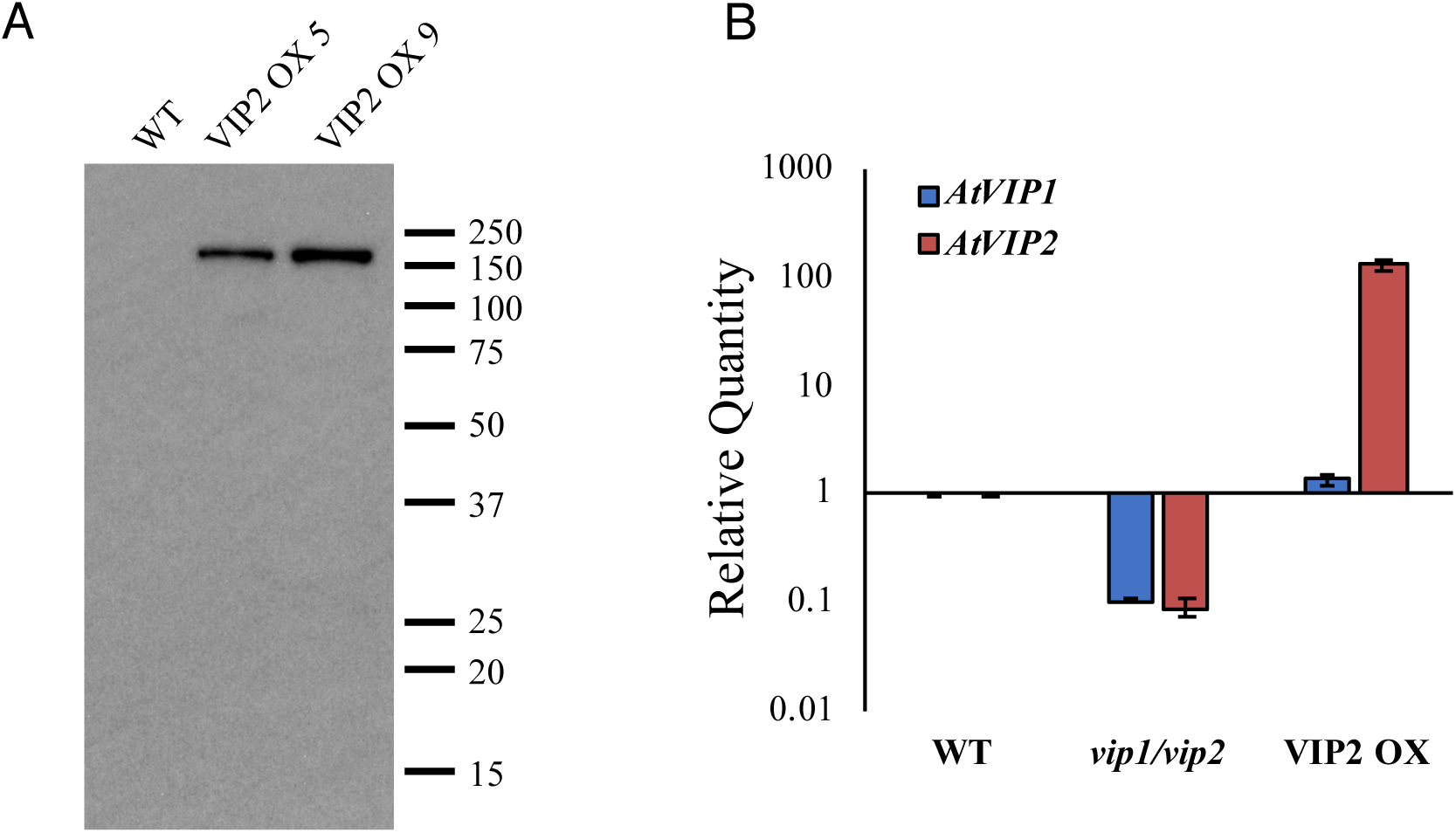
Characterization of VIP2 OX and vip1/vip2 double knock-out mutants. A, Immunoblot of GFP-Fusion Proteins in VIP2 overexpression (VIP2 OX) transgenic plant lines. 2 week-old seedlings from two different VIP2 OX lines were analyzed by immunoblotting using a mouse monoclonal anti-GFP antibody (Living Colors JL-8, Clontech). B, RNA preparation and qRT-PCR. RNA was isolated from 2 week-old seedlings of the indicated genotype and analyzed by quantitative RT-PCR using gene specific primers for *AtVip1* and *AtVip2* and SYBRgreen as described in Desai et al., 2014. Reactions were carried out in triplicate with PP2A as the reference gene. Relative expression was calculated by the ΔΔCT method for wildtype (WT), *vip1/vip2* double mutants and the VIP2 OX9 line.

**Supplemental Figure S5.**
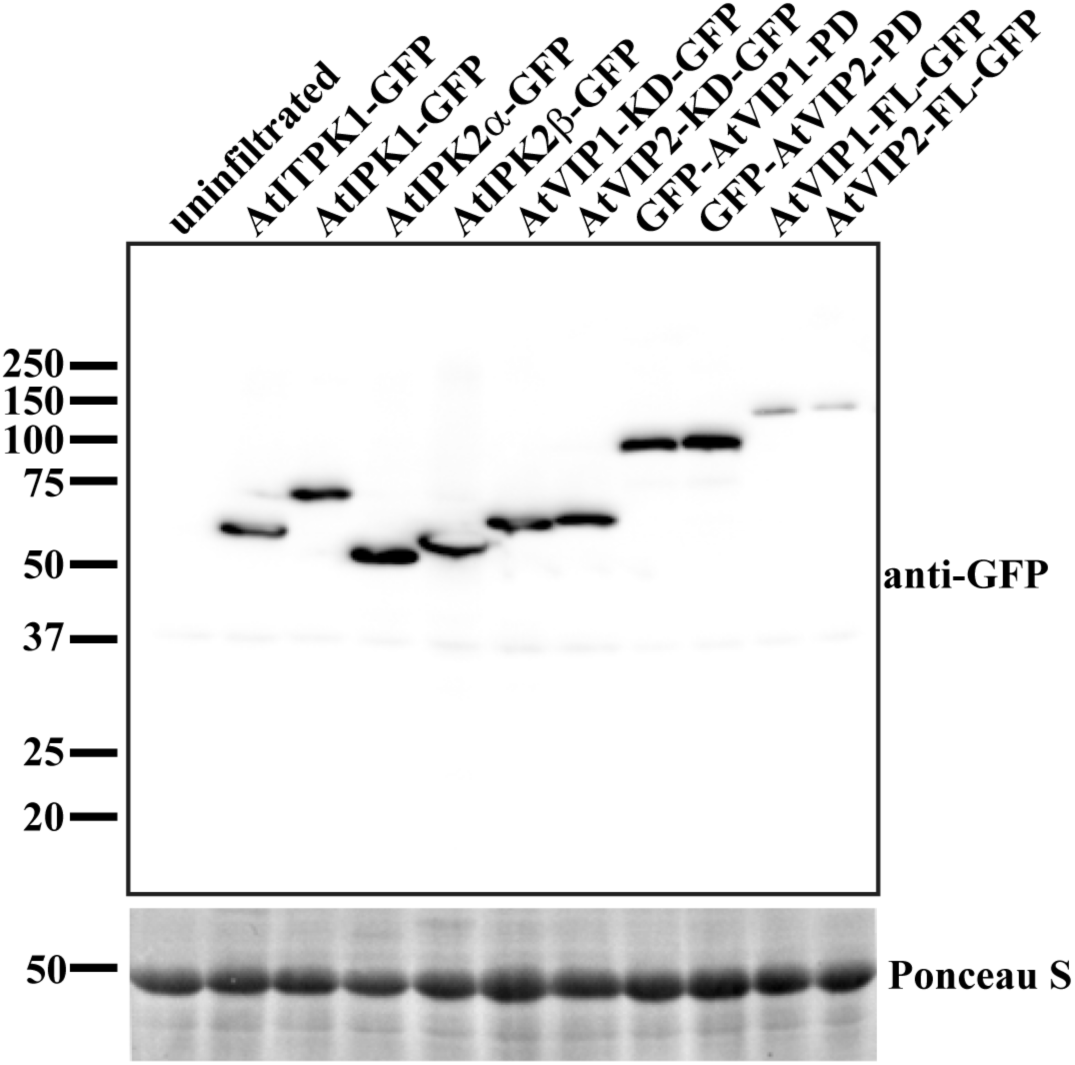
Immunoblot of GFP fusion proteins in infiltrated *N. benthamiani* leaves. *N. benthamiani* leaves were infiltrated with the indicated GFP fusion constructs, and 48 hours post-infiltration the leaf tissue was harvested and processed for immunoblot with anti-GFP. Ponceau S staining was used as a loading control.

**Supplemental Figure S6.**
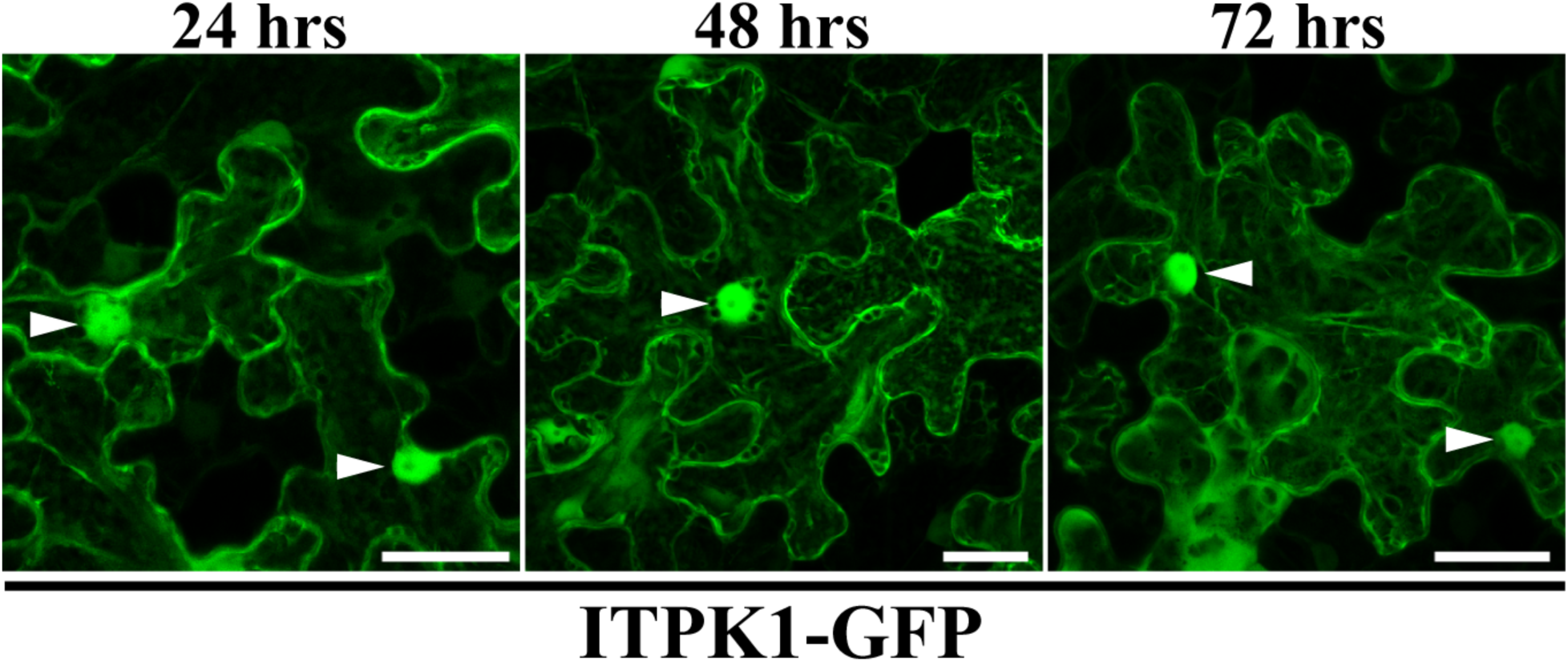
Time course of ITPK1-GFP expression. ITPK1-GFP was transiently expressed in infiltrated *N. benthamiana* leaves, and confocal Z-stacks were acquired at the indicated time points post-infiltration. Z-stacks are presented as maximum intensity projections. Arrowheads mark nuclei. Scale bars = 30 µm.

**Supplemental Figure S7.**
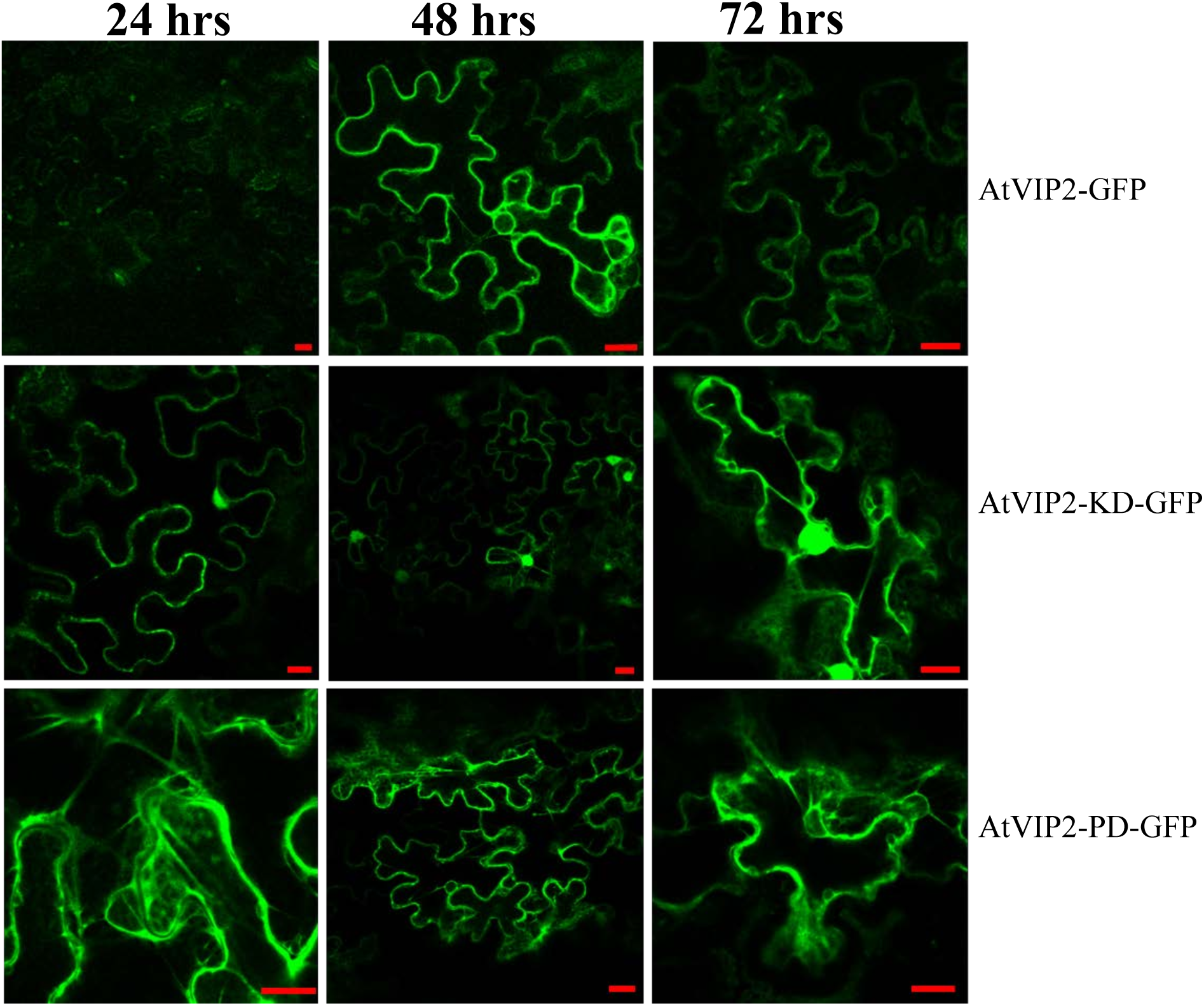
Time Course of VIP2FL-, VIP2KD-, VIP2PD-GFP Expression. VIP2FL-, VIP2KD- and VIP2-PD-GFP fusion proteins were transiently expressed in *N. benthamiana*. Leaves were imaged at 24, 48, and 72 hours post infiltration using confocal microscopy. No signal was detected at 12 hours post infiltration. Scale bar = 20 µm.

**Supplemental Table 1.**
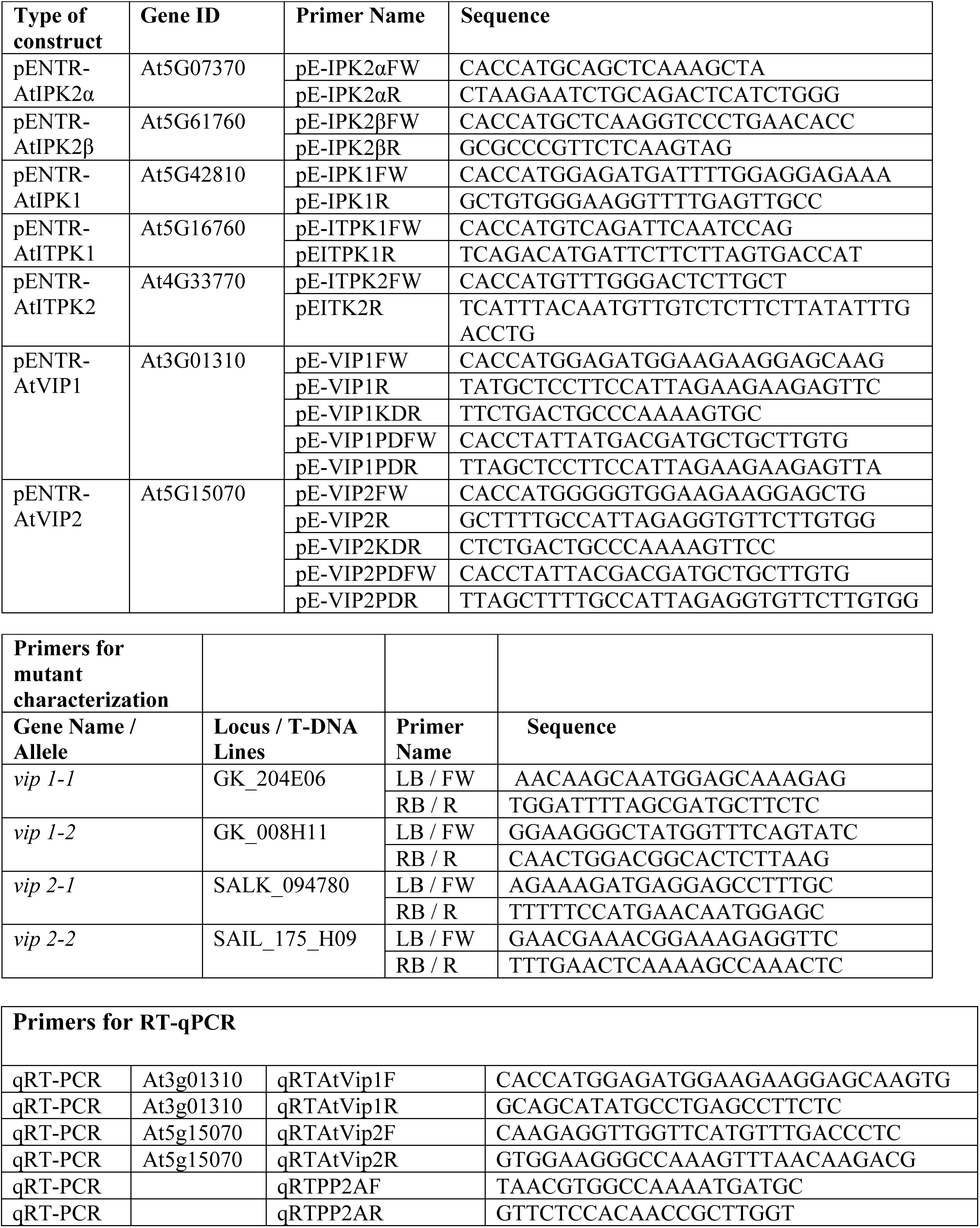
Oligonucleotide Primers Used in this Study.

